# Continual improvement of multiplex mutagenesis in Arabidopsis

**DOI:** 10.1101/2023.12.19.572307

**Authors:** Ward Develtere, Ward Decaestecker, Debbie Rombaut, Chantal Anders, Elke Clicque, Marnik Vuylsteke, Thomas B. Jacobs

## Abstract

CRISPR/Cas9 is currently the most powerful tool to generate mutations in plant genomes and more efficient tools are needed as the scale of experiments increases. In the model plant Arabidopsis, the choice of promoter driving Cas9 expression is critical to generate germline mutations. Several optimal promoters have been reported. However, it is unclear which promoter is ideal as they have not been thoroughly tested side-by-side. Furthermore, most plant vectors still use one of the two Cas9 nuclear localization sequence (NLS) configurations initially reported and can still be optimized. We genotyped more than 6,000 Arabidopsis T2 plants to test seven promoters and eleven NLS architectures across 14 targets to systematically improve the generation of single and multiplex inheritable mutations. We find that the RPS5A promoter and double-BP NLS architecture were individually the most efficient components. When combined, 99% of T2 plant contained at least one knockout mutation and 84% contained 4-7-plex knock-outs. These optimizations will be useful to generate higher-order knockouts in the germline of Arabidopsis and likely be applicable to other CRISPR systems as well.

## INTRODUCTION

Numerous studies have reported optimized CRISPR systems in *Arabidopsis thaliana* (Arabidopsis) and plants in general. CRISPR/Cas9 vectors can be optimized in multiple ways, including the promoter to express Cas9, the Cas9 nuclease sequence, the Cas9 nuclear localization signal (NLS), and the guide RNA (gRNA). Most studies in Arabidopsis have focused on comparing different promoters to generate single (simplex) or multiplex mutations (**Supplemental Table 1**). The cauliflower mosaic virus 35S promoter was initially used to drive Cas9 expression in Arabidopsis and rice^1^. However, it is now well established that the 35S promoter is not ideal to express Cas9 in Arabidopsis as it results in low insertion and deletion (indel) rates and almost no transmission to the germline. Other constitutive promoters such as UBQ1, UBI10, CsVMV, and especially RPS5A, have shown higher efficiencies^2–6^.

Tissue-specific promoters such as YAO, egg-cell (EC) 1.1, EC1.2, WOX2, DMC1, SPO11, P5, and NUC1 have been proposed to restrict Cas9 expression to the egg-cell and embryonic stage, which should result in early mutagenesis, thereby reducing somatic mutations and increasing the likelihood of identifying homozygous or bi-allelic mutations in the T1 generation^2–13^. Replacing the 35S promoter with an egg-cell specific promoter (EC1.1 or EC1.2) not only improved overall editing efficiency with up to 8% likely-triple mutants (determined through phenotyping) in T1 compared to 0% for 35S, but also reduced mosaicism in T1^11^. This was especially the case when combining the enhancer of *EC1.2* with the promoter of *EC1.1* (here, referred to as EC1) which led to 17% likely-triple mutants. While the EC1 promoter generated mutants with higher efficiency compared to 35S, it underperformed in the T2 generation compared to the constitutive RPS5A promoter, the tissue-specific YAO promoter and, in some cases, as compared to the UBI10 promoter^2^. In contrast, the efficiency of RPS5A was only slightly higher than EC1.2 in T2 (33% and 24%, respectively) and both promoters outperformed the constitutive PcUBI promoter (1%)^4^. Other tissue-specific promoters such as DMC1 and SPO11 generated only very few non-mosaic T2 plants (3.9% and 0.7%, respectively) compared to the YAO and meiosis-specific CDC45 promoters (24.5% and 22.7%, respectively)^3^. The latter also produced 3x more non-mosaic T1 mutants than the constitutive UBQ1 promoter when multiplexing 2-6 targets.

Despite the number of publications testing different promoters to drive Cas9 expression in Arabidopsis, due to inherent variations in methodologies across the studies (different gRNAs, simplex or multiplex, phenotyping or genotyping, T1 or T2 generation, different sample sizes etc.), it is difficult to make direct comparisons and determine the best promoter to produce germline mutations. Furthermore, almost all reports make comparisons at the T1 generation or in T2 plants which still carry the Cas9 transgene (**Supplemental Table 1**). Continuous activity by the CRISPR machinery in both of these materials makes it impossible to distinguish between germline and somatic mutations. In terms of multiplexing, only a single report has tested multiplex editing beyond two gRNAs^10^. As genome editing experiments become larger in scale and highly multiplex, as is the case with CRISPR screens^34, 35^, more efficient and thoroughly tested reagents are needed to reduce costs and increase the chances of success.

In comparison to the extensive optimization of Cas9 regulatory sequences in plants, the NLS architecture has received little attention. When Cas9 was first applied for mutagenesis in human and mouse cells, NLSs were fused to Cas9 to direct the prokaryotic protein to the eukaryotic nucleus. In one report, a human codon-optimized Cas9 with a single C-terminal SV40 NLS was used^14^. In a simultaneous report, a different human codon-optimized Cas9 was directed to the nucleus via an N-terminal SV40 NLS with a 3xFLAG tag and a C-terminal NLP NLS (SV40-NLP) and showed more efficient nuclear targeting than a single N-terminal SV40 NLS or C-terminal NLP NLS^15^. As CRISPR was rapidly adapted for plant genome editing, these two initial configurations (**Figure 1a**) have remained the basis of the vast majority of the plant vectors in use today, as exemplified by the most popular plant gene editing plasmids on Addgene (**Supplemental Table 2**).

**Figure 1:**
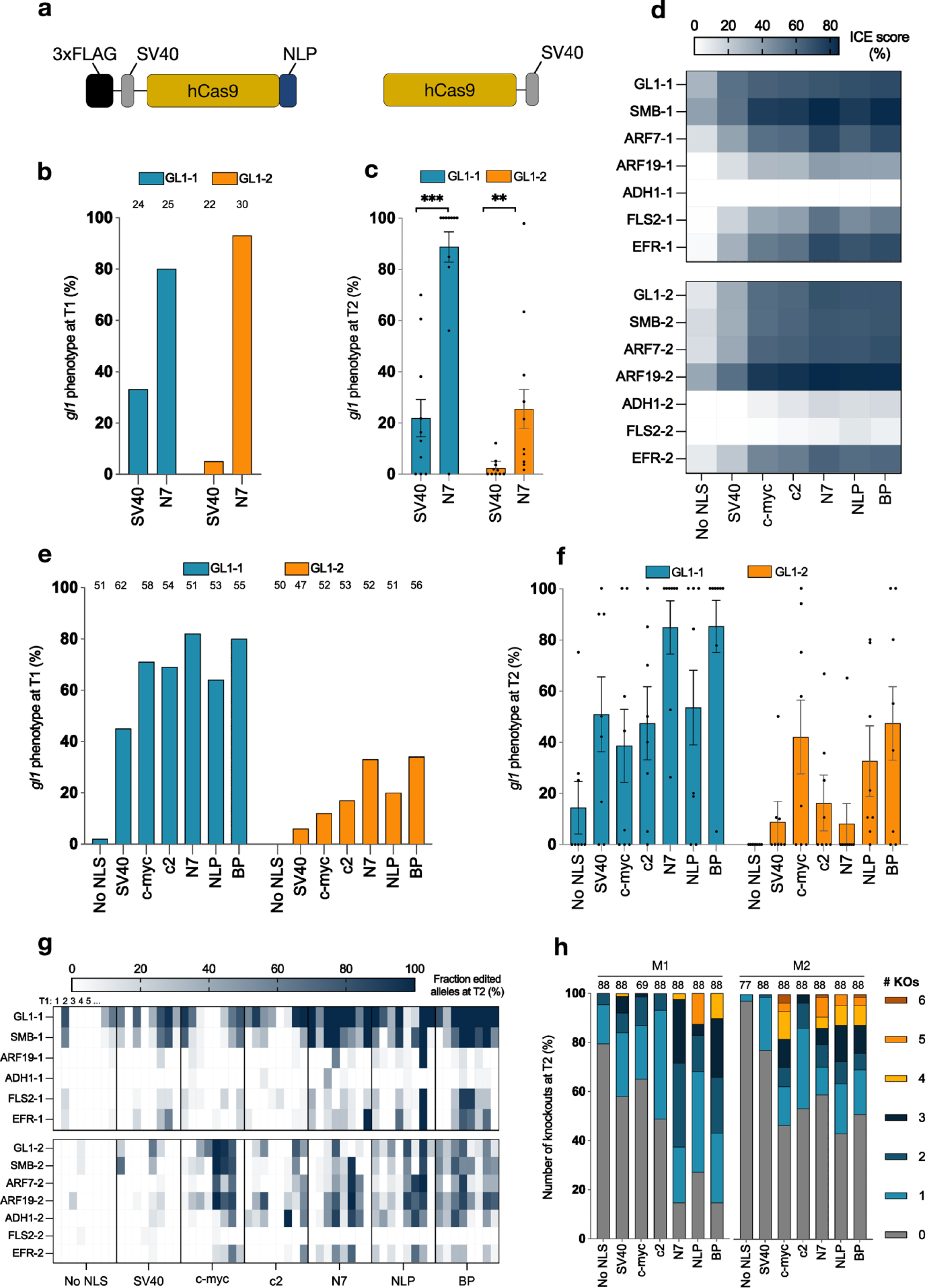
Editing efficiency of different C-terminal nuclear localisation signals for Cas9. (**a**) The two NLS architectures initially used in eukaryotic cells. Left: Cas9 with an N-terminal SV40 and a C-terminal NLP NLS^43^. Right: Cas9 with a C-terminal SV40 NLS^44^. (**b**) Percentage of T1 seedlings with a knockout phenotype for the GL1-1 and GL1-2 guide RNAs. Sample sizes are above the bars. (**c**) Phenotyping of Cas9-free T2 lines for the knockout phenotype (9 – 10 T1 lines per vector, 9 – 68 T2 seedlings per line). Bars represent the mean and error bars represent the standard error of the mean (Student’s two-sided t-test). (**d**) Average knockout efficiency (from three or four replicates) of different C-terminal NLSs targeting seven loci at a time (M1 top panel, M2 bottom panel) in PSB-D cell suspension cultures. The mean Inference of CRISPR Edits (ICE) score, which represents the proportion of cells with an indel, is given per target. (**e-f**) The percentage of knockout seedlings for both the GL1-1 and GL1-2 guide RNAs was determined at (**e)** T1 and (**f**) T2, respectively. 11 T1 lines per vector, 14 – 20 T2 seedlings per line. Bars represent the mean and error bars represent the standard error of the mean. No significant differences compared to the SV40 NLS were observed (Student’s two-sided *t*-test; p > 0.05) (**g**) Multiplex amplicon sequencing was performed on Cas9-free T2 plants. Each cell represents the fraction of edited alleles per line (8 T1 lines per vector, 5 – 11 T2 plants per line) per target (M1 top panel; M2 bottom panel). (**h**) The proportion of the number of knockouts per T2 plant (KO; homozygous or bi-allelic mutant) per vector. Only plants with genotyping data for at least 5 of the 6 or 7 targets were included for both M1 (left) and M2 (right). Sample sizes are above the bars. M1; GL1-1, SMB-1, ARF7-1, ARF19-1, ADH1-1, FLS2-1 EFR-1. M2; GL1-2, SMB-2, ARF7-2, ARF19-2, ADH1-2, FLS2-2, EFR-2. **, p < 0.01; ***, p < 0.001. Exact p-values are shown in Supplementary File 2.

Nuclear import is generally initiated by the formation of a ternary complex with importin α, importin β1 and the cargo^16^. There are two main types of NLSs: classical NLSs that are recognized by importin α and non-classical NLSs that bind directly to importin β family members^17^. Within the classical NLSs, there are six different classes, comprising classical monopartite (class 1 and 2), bipartite (class 6), minor site-specific (class 3 and 4) and plant-specific (class 5)^16^. Monopartite NLSs contain a single cluster of basic amino acids. Class 1 NLSs have at least four consecutive basic amino acids, KR(K/R)R or K(K/R)RK, as exemplified by the SV40 Large T-antigen NLS^18^. Class 2 NLSs have only three basic amino acids and are represented by K(K/R)X(K/R) as a putative consensus sequence^16^, such as the c-myc and c2 NLSs^19, 20^. Bipartite NLSs have two clusters of basic amino acids that are separated by a variable linker of 10-12 amino acids^16, 21^, such as the nucleoplasmin (NLP)^22^ and the bipartite SV40 (BP) NLSs^23^.

In human HEK293T cells, the addition of a linker between Cas9 and the N-terminal SV40 NLS increased nuclear targeting and mutagenesis efficiency and adding an N-terminal SV40 NLS to a Cas9 with a C-terminal NLP NLS (lacking the 3xFLAG tag) also increased activity^24, 25^. However, fusing an SV40 to either the N- or C-terminus, or to both ends did not alter the mutagenesis efficiency in zebrafish^26^. N- and C-terminal fusion of the BP NLS (double-BP) to Cas9 showed a 3-fold increase in nuclear targeting and a 1.5-fold increase in mutagenesis frequency as compared to the SV40 NLS in human embryonic stem cells and improved the BE4 base editor 1.3-fold in HEK293T cells^27, 28^. Not only is the type of NLS important, but also their configuration. The addition of both SV40 and NLP on the C-terminus showed improved nuclear targeting and 1.5- to 2-fold improvement in mutagenesis compared to different configurations with the same two NLSs^29^. In addition, increasing the number of either N- or C-terminal nuclear localization signals and replacing the common SV40 NLS with the c-myc NLS improved the knockout (KO) efficiency of AsCas12a^30^.

Systematic investigation of Cas NLS configurations has been limited in the plant field. In Arabidopsis, the removal of the C-terminal NLP NLS from the SV40-NLP configuration reduced mutagenesis below the background level of the assay^31^. It was shown that an SV40 NLS on both ends of a Cas9 containing 13 introns, instead of a single C-terminal SV40 NLS, increased the number of T1 plants with a KO phenotype from 58% to 72%^32^. In wheat and maize protoplasts, the double-BP architecture improved Cas9 and Cas12a adenine base editor (ABE) editing rates compared to a C-terminal triple SV40 NLS^33^.

Here, we systematically test different NLS and promoter configurations for the production of inheritable, multiplex mutants in Arabidopsis. Our standard vector architecture was based on pDE-Cas9, in which the C-terminal SV40 configuration was codon-optimized for Arabidopsis and expressed under the control of the *Petroselinum crispum* UBIQUITIN 4-2 (PcUBI) promoter^36^, although we use a different vector backbone and the G7 terminator (G7T) instead of the pea3A terminator. Using this as our standard, we systematically tested up to seven promoters and six NLSs (in different architectures) for both simplex and multiplex editing. We performed our comparisons on the transgene-free T2 generation as it allows us to unequivocally quantify germline edits which would be the foundational generation for most users. We find that the combination of the RPS5A promoter with the double-BP NLS architecture leads to the most multiplex-edited plants, with 99% of T2 plants containing at least one KO mutation and >80% of plants contain 4-7-plex KOs. These vectors will be useful when creating highly multiplex KO lines in Arabidopsis.

## RESULTS

### The N7 NLS is an efficient nuclear localization signal and increases Cas9 mutagenesis efficiency

The N7 NLS was previously identified from a GFP-tagged cDNA library in Arabidopsis. This library generated a line (N7) containing the C-terminal part of the Ankyrin repeat family protein (AT4G19150) in which the GFP was targeted to the nucleus^37^. The C-terminus was later combined with a linker to construct the pGGD007 cloning module in the GreenGate system^38^. Analysis of the linker-N7 NLS sequence with cNLS Mapper^39^ predicts one monopartite NLS and two bipartite NLSs (**Supplemental Figure 1a**). As the linker-NLS was considerably longer than other NLSs and there was no predicted NLS in the first part of this sequence, we removed the (GS)_7_-G linker sequence and created a novel Golden Gate entry module, pGG-D-NLS_N7-E (**Supplemental Figure 1a**). The N7 NLS showed very efficient nuclear targeting of GFP in tobacco leaf-infiltration experiments, especially compared to SV40 (**Supplemental Figure 1b**).

Considering that SV40 resulted in poor GFP nuclear localization and that a single C-terminal SV40 was our standard for Cas9 experiments, we wondered if Cas9 mutagenesis efficiency could be improved by modifying the NLS. We set up a pilot experiment to test this by comparing SV40 and N7 Cas9 mutagenesis efficiency in Arabidopsis. We expressed GFP-Cas9-SV40 and GFP-Cas9-N7 under the control of the PcUBI promoter and G7 terminator (G7T). We targeted the Arabidopsis *GLABRA1* (*GL1; AT3G27920*) gene with two gRNAs (GL1-1 or GL1-2) separately. *GL1* is required for the formation of trichomes, so targeting it provides a simple phenotype as KO plants lack trichomes on the leaves and stems^40^.

The four GFP-Cas9-SV40 and GFP-Cas9-N7 vectors were transformed into Arabidopsis using floral dip^41^ and transgenic events selected with a fluorescent accumulating seed technology (FAST) system^42^. Fourteen days after sowing, T1 seedlings were screened for KO phenotypes. Plants completely lacking trichomes were categorized as KOs, whereas plants showing a mosaic phenotype (somatic mutagenesis) were categorized as wild-type (WT). For both gRNAs, GFP-Cas9-N7 showed 2-18 fold more KO T1 plants than GFP-Cas9-SV40 (**Figure 1b**).

As we cannot distinguish between somatic or germline mutations in the T1 generation, we analyzed the transmission of the mutations in transgene-free (null segregant) T2 plants. Approximately 50 FAST-negative T2 seeds from 10 T1 lines were phenotyped per vector. As anticipated, mosaic plants were not observed, indicating that the Cas9 system was absent. For both targets, the N7 vector generated 4-14 fold more KO plants than SV40 (**Figure 1c**). Genotyping results from 38 FAST-negative T2 plants from two to three parental T1 lines per vector confirmed the presence of homozygous and bi-allelic mutations in the *GL1* gene (**Supplemental Table 3**).

### Screening different Cas9-NLS fusions for multiplex mutagenesis

These results confirmed that the mutation frequency could be increased by modifying the NLS. We then wondered which NLS would lead to the highest level of mutagenesis. To determine this, we performed a screen with six NLSs fused to the C-terminus of Cas9. We selected a panel of NLSs to cover the range of NLS classes and ones routinely used in the plant and gene-editing fields: SV40, c-myc, c2, N7, NLP, and BP. A no-NLS control was included as a negative control. We targeted seven genes in multiplex: *GL1*, *SMB* (AT1G79580), *ARF7* (AT5G20730), *ARF19* (AT1G19220), *ADH1* (AT1G77120), *FLS2* (AT5G46330) and *EFR* (AT5G20480) using gRNAs based on the reference genome sequence of Arabidopsis Col-0. The GL1-1, SMB-1, SMB-2 ARF7-1, ARF7-2, ARF19-1, and ARF19-2 gRNAs were previously reported^40, 42^, but none of the gRNAs were selected based on predicted or known efficiency. Two gRNA arrays (referred to as M1 and M2) were constructed with each array containing one gRNA per gene. Each gRNA was expressed from an individual transcriptional unit by the *AtU6-26* promoter. A separate mCherry-NLS cassette was included for fluorescence-activated cell sorting (FACS)^45^.

The multiplex mutagenesis vectors were transformed into PSB-D cell suspension cultures. At 17 days post-cocultivation, the cultures were protoplasted and sorted for mCherry-positive protoplasts by FACS. DNA was extracted from the sorted samples and genotyped via ICE^46^. As the gRNAs were designed to target the Arabidopsis Col-0 reference genome, three gRNAs (ARF7-1, ADH1-1 and ADH1-2) had mismatches to the target sites in the PSB-D cell suspension culture, which is derived from the Landsberg *erecta* genotype. All target sites show a range of mutagenesis from ∼5-80%, except the ADH1-1 target which was non-functional (**Figure 1d**). This target has a mismatch within the gRNA seed region where mutations are more likely to disrupt target recognition by Cas9^47, 48^. The vectors without an NLS sequence showed the lowest indel efficiencies with an average of 12% across all targets (**Supplemental Figure 2a**). Mutagenesis was improved with the addition of the SV40 NLS (28%), however, it produced significantly fewer indels compared to all other NLSs. The highest indel efficiencies were obtained with N7 (54%), NLP (52%) and BP (54%).

The fourteen vectors were also transformed into Arabidopsis plants via floral dip. Around 50 FAST-positive T1 seeds per vector were sown *in vitro* and phenotyped for the presence or absence of trichomes two weeks after sowing (**Figure 1e**). Consistent with previous results, GL1-1 shows a higher number of KO plants than GL1-2. The No-NLS vectors had 2% and 0% KO plants for the GL1-1 and GL1-2 targets, respectively. These numbers were improved to 45% and 6% with the SV40 NLS, but again, SV40 was inferior to all other NLSs tested. N7 and BP had the highest number of T1 KO plants with ∼80% for GL1-1 and ∼30% for GL1-2.

To check for inheritance, 11 FAST-negative T2 seeds were selected from eight independent T1 lines for each vector and sown *in vitro*. Two weeks after sowing, plants were phenotyped, harvested for DNA extraction and genotyped using multiplex amplicon sequencing (MAS). Illumina sequencing of PCR products followed by SMAP-haplotype window analysis^49^ was used to genotype each of the seven target sites in each plant. Across all samples, indel frequencies had clear peaks at 0, 50, and 100% as expected for a diploid (**Supplemental Figure 3**). We set indel score thresholds of <20% as wild-type, 20-80% as heterozygous and >80% as KO. Knockout mutations were further subdivided into homozygous (two identical mutant alleles) or bi-allelic (two distinct mutant alleles). We also scored genotypes as “false” if we observed 0 or >2 alleles for each target site in each plant. The ARF7-1 target site resulted in only false calls and was removed from the analysis. From a total of 8,624 possible genotype calls (14 vectors x 8 T1 lines x 11 T2 plants x 7 targets), we were able to capture information for 8,056 (93%) and had near complete coverage with six or seven genotype calls for 1,166 plants (**Supplemental Figure 4**).

The T2 phenotyping largely mirrored those of the T1; for GL1-1, N7 and BP led to the highest average number of KO plants (85%) and all lines contained at least one KO plant (**Figure 1f**) although the results were not significantly different than SV40. The results for GL1-2 were lower and more variable; c-myc, NLP and BP performed the best with 32-47% of the T2 plants containing *GL1* KOs. The T2 phenotyping results correlated well with the genotyping (**Figure 1g-h**). Across all targets, N7, BP, and NLP had a significantly higher fraction of edited alleles than SV40 (**Supplemental Figure 2b**). Overall, there are clear gRNA and T1-line effects (**Figure 1g**). FLS2-2 and ADH1-1 were the least active gRNAs, consistent with the results from PSB-D cell cultures. One No-NLS M1 T1 line had indels in five of the six target sites and was responsible for most of the editing observed with that vector (**Figure 1g**), with 20% of the T2 plants containing simplex and duplex KO mutations (**Figure 1h**). When looking at the number of KO mutations per vector, NLP had the highest number of multiplex KOs for M1, but this is largely attributed to a single line (**Figure 1g-h**). Across both gRNA arrays, N7, NLP and BP had the highest number of multiplex KOs. We also looked at the type of DNA repair products and, as expected for Cas9 mutagenesis, we observed primarily +1 insertions and there are no obvious differences between the different NLSs (**Supplemental Figure 5**).

### Further increasing multiplex editing efficiency by combining NLSs

As our NLS screen was performed with only C-terminal fusions to Cas9, we wanted to test if combining multiple NLSs would further increase editing efficiency. We were unsure if there would be specific N- and/or C-terminal effects, so we performed a test of the N7 and BP NLSs in the four possible combinations with the M1 and M2 arrays (**Figure 2a**). We included the C-terminal SV40 and SV40-NLP architectures as these are standards in the field, and the C-terminal BP and N7 from the first screen as controls. These 16 vectors were stably transformed into Arabidopsis via floral dip and FAST-negative T2 plants from eight T1 lines were evaluated via phenotyping and MAS. Phenotyping showed similar results as the C-terminal fusions. We observed significantly more *GL1* KOs with the C-terminal BP and double-BP architectures as compared to SV40 (**Figure 2b**). Interestingly, the SV40-NLP architecture produced 82% T2 KOs with GL1-1 but nearly zero for GL1-2. The double-BP architecture was by far the most efficient with every T1 line containing T2 KOs and an average of >90% for both gRNAs.

**Figure 2:**
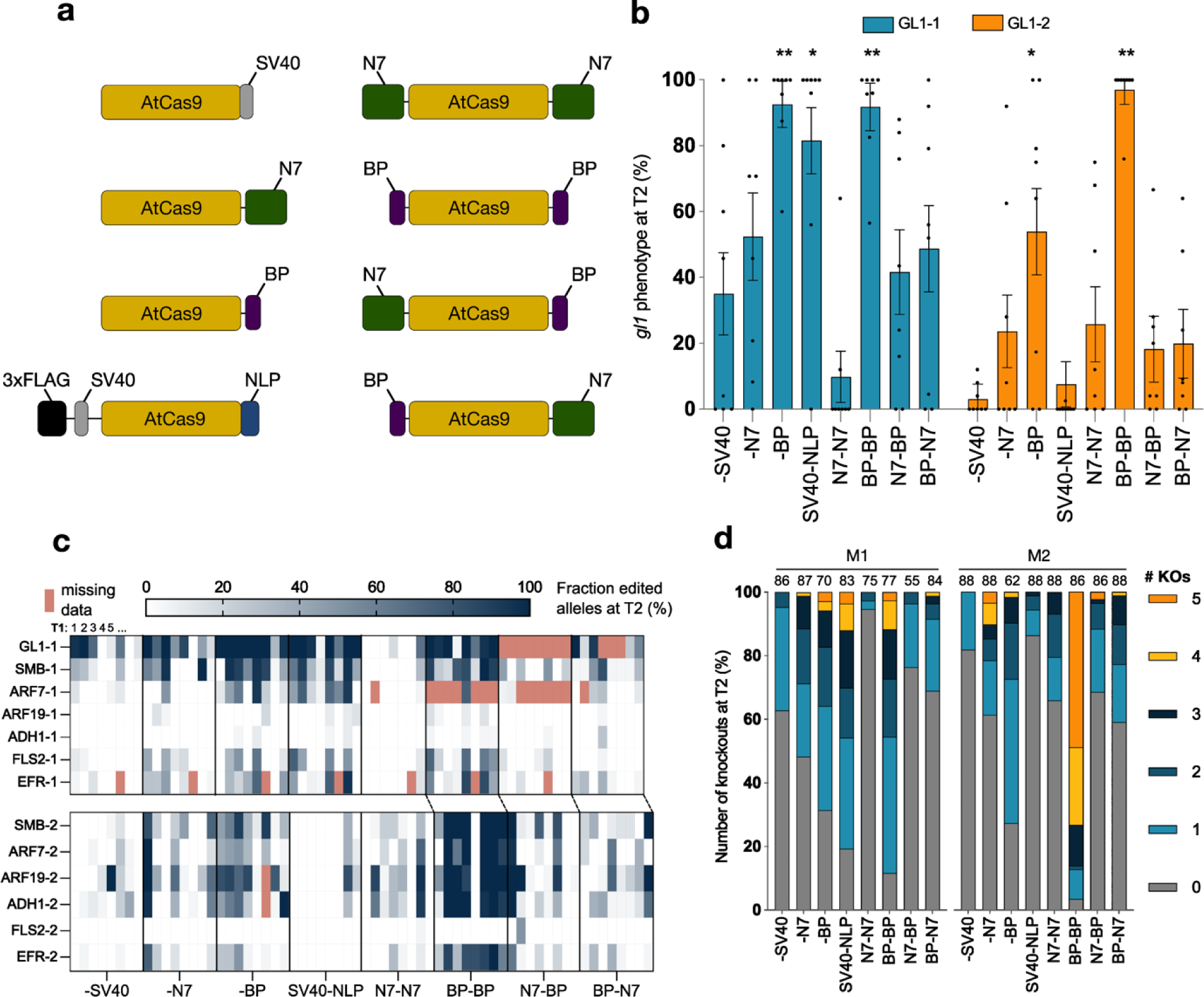
Editing efficiency of different combinations of BP and N7 NLS Cas9 architectures. (**a**) Schematic of the eight different NLS architectures tested. (**b**) Phenotyping of Cas9-free T2 lines for a knockout phenotype (8 T1 lines per vector, 22 – 25 T2 plants per line). Bars represent the mean and error bars represent the standard error of the mean. Statistical significances represent comparisons of each architecture with the -SV40 architecture (Student’s two-sided *t*-test). (**c**) Multiplex amplicon sequencing results for the different NLS architectures (7 - 8 T1 lines per vector, up to 11 T2 plants per line). Each cell represents the fraction of edited alleles per line per target (M1 top panel; M2 bottom panel). Lines with fewer than 5 genotyped plants at a particular target site were excluded (missing data; red cells). (**d**) The proportion of the number of knockouts per T2 plant (KO; homozygous or bi-allelic mutant) per vector. Only plants with genotyping data for at least 5 of the 6 or 7 targets were included for both M1 (left) and M2 (right). Sample sizes are above the bars. M1; GL1-1, SMB-1, ARF7-1, ARF19-1, ADH1-1, FLS2-1 EFR-1. M2; GL1-2, SMB-2, ARF7-2, ARF19-2, ADH1-2, FLS2-2, EFR-2. *, p < 0.05; **, p < 0.01. Exact p-values are shown in Supplementary File 2.

Unfortunately, the GL1-2 amplicon failed in this MAS experiment and was removed from the analysis (**Figure 2c**). Across all targets, N7, BP, SV40-NLP and double BP performed significantly better than SV40 (**Supplemental Figure 6**). By far, the double-BP architecture performed the best of all tested with >88% of all T2s containing at least one KO gene and 49% of T2 plants containing 5-plex KOs for M2 (out of 6 possible; **Figure 2d**). As before, the DNA repair products were primarily 1-bp insertions (**Supplemental Figure 7**).

The combinations of the BP and N7 NLSs gave surprisingly negative results and were equivalent to SV40 (**Supplemental Figure 6**). With the double-N7 vector, only 5% of the plants contained KO mutations in the M1 array. This was worse than the single C-terminal N7 architecture (52%) and even worse than SV40 (47%) for the M1 array. Both the BP-N7 and N7-BP architectures induced fewer KOs than the C-terminal BP, suggesting that these two NLSs do not combine well. While we are missing genotyping data for the GL1-2 targets and GL1-1 for N7-BP, the phenotyping results confirm the average-to-low efficiency of those combinations (**Figure 2b**).

### Evaluation of seven promoters to express Cas9

While the NLS experiments were ongoing, we also performed a promoter screen with some of the top Arabidopsis Cas9 promoters reported in the field: PcUBI (our lab standard), RPS5A, EC1, and YAO plus the CLV3, HMG and P16 promoters that had previously shown high rates of germline editing in Arabidopsis using Cre/lox and/or TALENs^50, 51^. We used a single C-terminal SV40 NLS on Cas9 as this was our lab standard and also a P2A-mCherry-N7 at the end of the Cas9 expression cassette^42^ (**Figure 3a**). Fourteen expression vectors (7 promoters x 2 *GL1* gRNAs) were transformed into Arabidopsis via floral dip, FAST-positive T1 seeds were selected and the seedlings were phenotyped two weeks after sowing. The GL1-1 gRNA resulted in KO phenotypes with all promoters whereas GL1-2 failed to produce any KOs with the YAO and CLV3 promoters (**Figure 3b**). The PcUBI and RPS5A promoters were the most consistent, with >60% of T1s having a KO phenotype.

**Figure 3:**
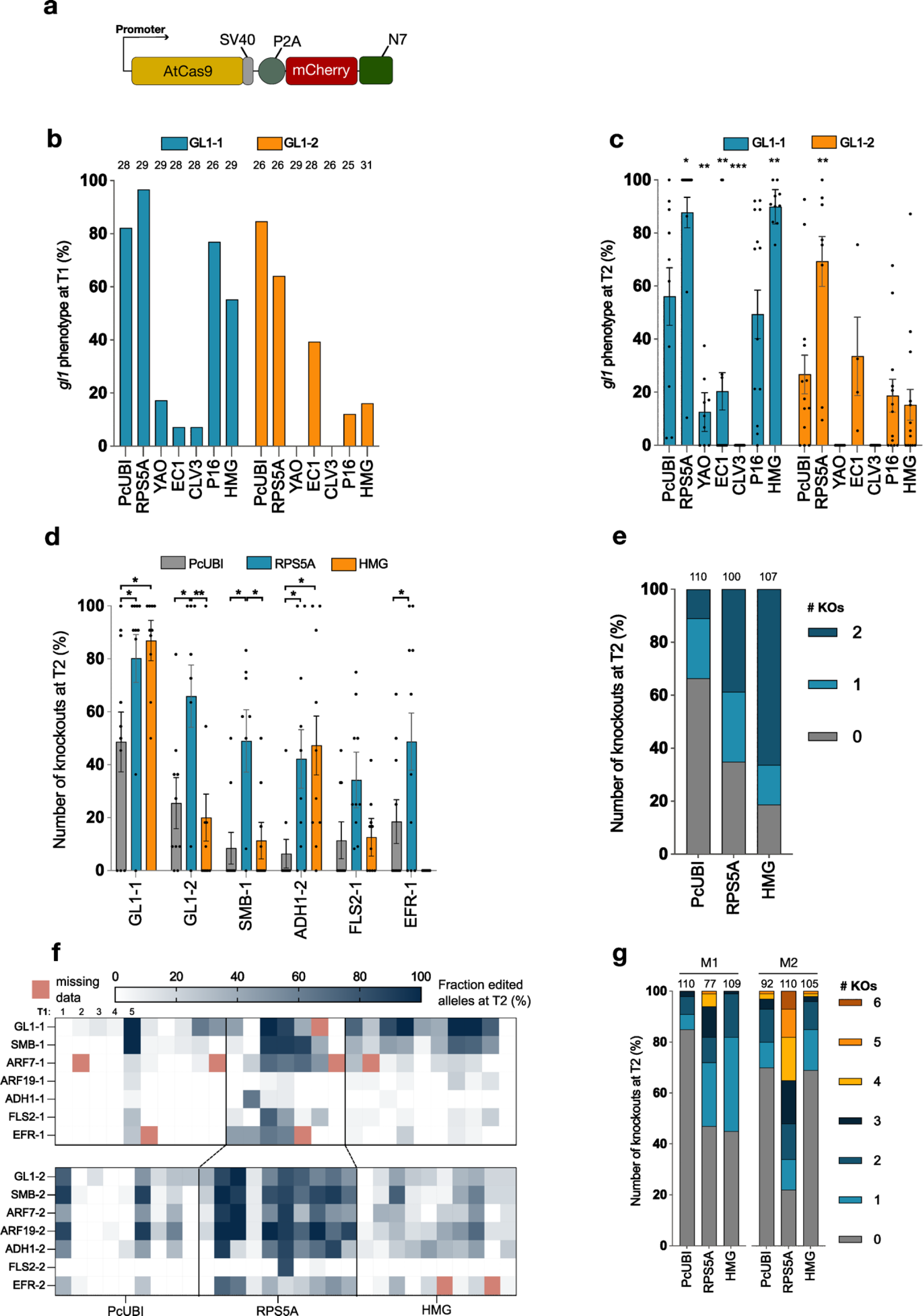
Editing efficiency of different promoters to express Cas9. (**a**) Vector architecture used to compare different promoters to express Cas9. Cas9 contains a C-terminal SV40 NLS followed by a P2A ribosomal skipping peptide and an mCherry cassette with a C-terminal N7 NLS. (**b**) The percentage of T1 seedlings with a knockout phenotype for the GL1-1 and GL1-2 guide RNAs per promoter. Sample sizes are above the bars. (**c**) Phenotyping of Cas9-free T2 lines for a knockout phenotype (4 - 13 T1 lines per vector, 19 – 70 T2 seedlings per line). Statistical significances represent comparisons of each promoter with the PcUBI promoter (Student’s two-sided *t*-test). (**d**) Multiplex amplicon sequencing results for the additional targeted loci in simplex (8 – 10 T1 lines per vector, up to 11 T2 plants per line). Bars represent the mean number of knockouts and error bars represent the standard error of the mean (Student’s two-sided *t*-test). (**e**) Multiplex amplicon sequencing results for vectors targeting ARF7-2 and ARF19-2 in duplex (10 T1 lines per vector, up to 11 T2 plants per line). The proportion of the number of knockouts per T2 plant (KO; homozygous or bi-allelic mutant) per vector is given. (**f**) Multiplex amplicon sequencing results for vectors targeting M1 and M2 in multiplex (7 – 10 T1 lines per vector, up to 11 T2 plants per line). Each cell represents the fraction of edited alleles per line per target (M1 top panel; M2 bottom panel). Lines with fewer than 5 genotyped plants at a particular target site were excluded (missing data; red cells). (**g**) The proportion of the number of knockouts per T2 plant (KO; homozygous or bi-allelic mutant) per vector. Only plants with genotyping data for at least 5 of the 7 targets were included for both M1 (left) and M2 (right). Sample sizes are above the bars. M1; GL1-1, SMB-1, ARF7-1, ARF19-1, ADH1-1, FLS2-1 EFR-1. M2; GL1-2, SMB-2, ARF7-2, ARF19-2, ADH1-2, FLS2-2, EFR-2. *, p < 0.05; **, p < 0.01; ***, p < 0.001. Exact p-values are shown in Supplementary File 2.

Per vector, up to 13 FAST-positive T1 seeds were propagated and per T1 line, 31-70 FAST-negative T2 seeds were phenotyped (**Figure 3c**). The RPS5A promoter was the most efficient with an average KO efficiency of 89% and 66% for both *GL1* gRNAs, respectively, and every line produced at least one KO T2 seedling. HMG was equally effective with GL1-1, but equivalent to the other promoters with GL1-2. The CLV3 promoter was completely ineffective as it did not produce any KOs at T2.

The EC1 promoter was surprisingly one of the worst for GL1-1 with, on average, only 19% of T2s having KO mutations, but it did produce two lines with 100% KO at T2. For GL1-2, EC1 was similar to the other promoters (except RPS5A) but we could only evaluate four lines due to a large number of escapes (62% of the lines). As the RbcS terminator was shown to be important for the EC1 promoter activity^11^ and we used the G7 terminator, we decided to repeat the experiment for EC1 and also test the replacement of the G7 terminator with RbcS. The RPS5A and HMG promoters were included for comparison. However, in this experiment our original vector with the EC1 promoter and G7 terminator was completely ineffective at the GL1-2 target site (**Supplemental Figure 8a**). By comparison, the results from the RPS5A and HMG promoters were equivalent to the first promoter experiment. Based on these results, we conclude that the EC1 promoter performs inconsistently under our conditions and therefore continued with only PcUBI, RPS5A, and HMG.

We next tested if these three promoters could be generalized to more targets in simplex and multiplex. We cloned gRNAs targeting *ADH1*, *SMB*, *EFR*, and *FLS2* in simplex and *ARF7* and *ARF19* in duplex. Per vector, we genotyped 11 FAST-negative T2s from 10 T1 lines via MAS. Overall, RPS5A resulted in the most consistent number of KO plants per line in T2, with 35-80% containing KOs (**Figure 3d**). HMG was particularly effective for GL1-1 and ADH1-2, however, it was no different than RPS5A and there were essentially no KOs at EFR-1. In duplex, HMG was significantly better than PcUBI, with 81% of plants having a KO for at least one gene and 66% of T2s with a double KO (**Figure 3e, Supplemental Figure 8b**).

In the multiplex experiments, the RPS5A promoter was significantly better than PcUBI and HMG and resulted in the most consistent KO induction with 44% of all alleles edited (**Figure 3f-g, Supplemental Figure 8c**). For the M1 array, RPS5A and HMG produced approximately the same number of KO plants, but there are even higher-order mutants for RPS5A due to the increased editing at ARF7-1, FLS2-1, and EFR-1 (**Figure 3f-g**). For the M2 array, RPS5A clearly produced the most KOs and 7% of the T2 plants contained KOs in six of the seven target genes. T1 line effects are also apparent. The FLS2-2 gRNA is the least efficient in the arrays, but a single RPS5A line produced all of the KOs for this target and just three T1 lines are responsible for most of the KOs for PcUBI with the M1 and M2 arrays (**Figure 3f**). The DNA repair profiles are also similar for the different promoters (**Supplemental Figure 9**). Taken together, it is clear that the RPS5A promoter is the most efficient and consistent at producing multiplex KO plants in Arabidopsis.

### Combining the double-BP architecture and RPS5A promoter results in highly multiplex KO plants with few off-targets

Since the NLS screen was performed with the PcUBI promoter and the promoter screen with the SV40 NLS, our two standards at the time, an obvious experiment was to test the combination of the double-BP NLS architecture with the RPS5A promoter (**Figure 4a**). For this we relied on the same multiplex arrays, M1 and M2, and included the best vectors (PcUBI::BP-BP and RPS5A::-SV40-P2A-mCherry-N7) from the previous tests as side-by-side controls. As before, we phenotyped 11-22 FAST-negative T2s from 10 T1 lines two weeks after sowing. The results were similar to the earlier experiments, except RPS5A::-SV40-P2A-mCherry-N7 resulted in only an average of 34% of KO T2 plants with GL1-2 (**Figure 4b**). The RPS5A::BP-BP vector resulted in a similar number of *GL1* KOs as the PcUBI::BP-BP, with both having on average >90% KO at T2s.

**Figure 4:**
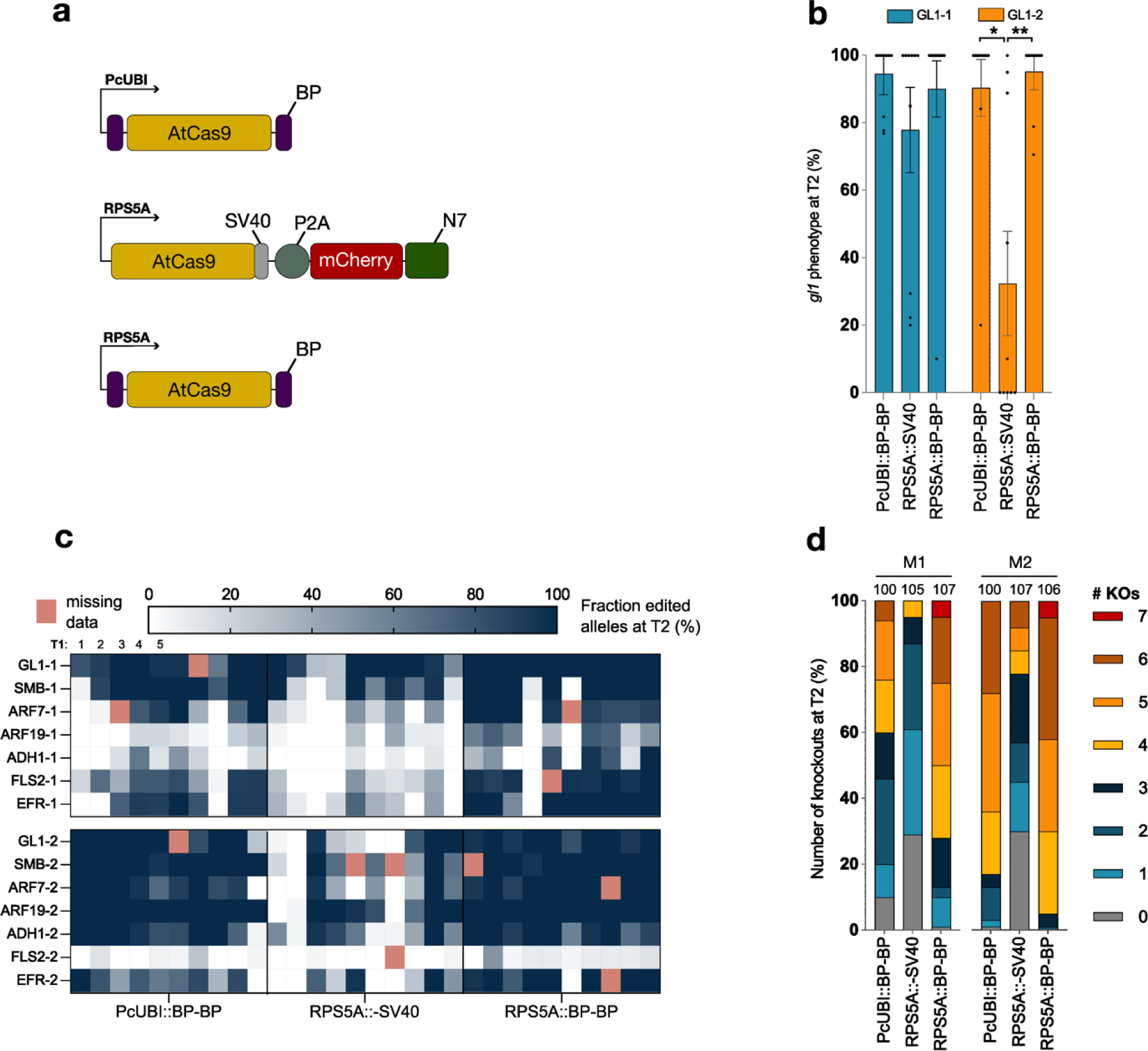
Editing efficiency of vectors combining promoter and NLS optimizations. (**a**) The schematics of the two top-performing architectures from previous experiments and a combination of both architectures. (**b**) Phenotyping of Cas9-free T2 lines for a knockout phenotype at the GL1-1 and GL1-2 target sites (10 T1 lines per vector, 11 – 20 T2 plants per line). Bars represent the mean number of knockouts and error bars represent the standard error of the mean (Student’s two-sided *t*-test). (**c**) Multiplex amplicon sequencing results for the vectors targeting M1 and M2 in multiplex (10 T1 lines per vector, up to 11 T2 plants per line). Each cell represents the fraction of edited alleles per line per target (M1 top panel; M2 bottom panel). Lines with fewer than 5 genotyped plants at a particular target site were excluded (missing data, red cells). (**d**) The proportion of the number of knockouts per T2 plant (KO; homozygous or bi-allelic mutant) per vector. Only plants with genotyping data for at least 5 of the 7 targets were included for both M1 (left) and M2 (right). Sample sizes are above the bars. M1; GL1-1, SMB-1, ARF7-1, ARF19-1, ADH1-1, FLS2-1 EFR-1. M2; GL1-2, SMB-2, ARF7-2, ARF19-2, ADH1-2, FLS2-2, EFR-2. *, p < 0.05; **, p < 0.01. Exact p-values are shown in Supplementary File 2.

Per vector, we genotyped 11 FAST-negative T2s from 10 T1 lines via MAS. The combination of the RPS5A promoter and double-BP architecture led to significantly more edited alleles than the other two vectors, with an average of 78% of all alleles edited and 99% of all plants containing at least one KO mutation (**Figure 4c-d, Supplemental Figure 10**). From both arrays, 5% of the T2 plants contained 7-plex KOs, largely due to increased activity at inefficient gRNAs (e.g. ARF19-1 and ADH1-1; **Figure 4c**). We observe largely the same DNA repair products for all vectors, though RPS5A::BP-BP produced the fewest number of 1-bp insertions for 11 of the 14 target sites (**Supplemental Figure 11**). Together, these results clearly show that RPS5A::BP-BP is our best architecture at inducing multiplex, inheritable KOs in Arabidopsis.

As the rate of mutagenesis increases, the concern for off-target mutations also increases. The GL1-1 gRNA has multiple putative off-target sites that differ by only one or two nucleotides as determined by CCTop^52^. To check if the increased on-target activity using the RPS5A::BP-BP vector would lead to a greater number of off-targets, we genotyped six T2 plants from ± six T1 lines at four GL1-1 off-target sites (*MYB62*, *MYB114*, *RAX2* and *MYB60*) using Sanger sequencing. From 149 genotype calls, we only identified two off-target events. One heterozygous mutation at *MYB60* and a homozygous AA insertion at *MYB114* (**Supplemental Figure 12**). Thus, despite the significant increase in on-target editing rates, we observe minor off-target activity with the GL1-1 gRNA.

One recurring question during the optimization campaign was when in the T1 plant development were the germline mutations being made in the more efficient vectors. Previous research has suggested that mutations arising early in the germline would decrease the allelic diversity^11^. To investigate this in our own experiments, we looked at the diversity of alleles in each line by counting the number of unique T2 alleles that were produced from each T1 line. We divided this by the editing efficiency for each particular gRNA per line to account for variations between gRNAs and lines (**Supplemental Figure 13**). Overall, we see that the most efficient gRNAs (e.g. GL1-1 and SMB-1) have the lowest allelic diversity per line across all experiments and that our least efficient gRNAs (e.g. ARF19-1 and ADH1-1), when they do produce germline mutations, result in the greatest allelic diversity per line. Similarly, the most efficient vectors (e.g. PcUBI::BP-BP and RPS5A::BP-BP) produce very little allelic diversity at almost all targets. The final RPS5A::BP-BP produced, on average, 3 - 4 unique alleles per target per line (**Supplemental Figure 13c**). These data suggest that both the promoter and NLS optimizations shift mutations to earlier timepoints during plant development as compared to the less efficient vectors.

## DISCUSSION

Numerous optimizations to CRISPR/Cas tools have been reported in plants. Yet further improvement is essential as researchers continue to increase the scale of experiments using multiplexing. In this work, we show that simplex and multiplex CRISPR/Cas9 mutagenesis efficiency in Arabidopsis can be significantly increased over the two predominant NLS architectures by using a double-BP NLS, where the BP NLS is attached to both the N- and C-termini of Cas9. Similarly, we demonstrate that the RPS5A promoter to express Cas9 is the most consistent and efficient at inducing inheritable, multiplex KOs in Arabidopsis.

An increase in editing efficiency is particularly important when performing CRISPR screens to keep the mutant population small^34, 35, 53^. For example, when attempting to capture all 190 double mutants in a pool of 20 genes, a CRISPR vector with two gRNAs per vector and an overall mutation efficiency of 31% (e.g. RPS5A::Cas9-SV40-P2A-mCherry-N7) will require an estimated population size of ∼18,000 individuals^53^. A vector that confers 73% global mutation efficiency (e.g. RPS5A::BP-Cas9-BP) would reduce this population to just ∼3,000. This represents a considerable cost and time savings even for a relatively small plant like Arabidopsis.

We screened different NLS classes and consistently find that the class 6 bipartite NLSs lead to the highest levels of germline editing. Further improvements may be possible by including more bipartite NLSs or, for instance, making specific combinations with BP and NLP. Our results and others show that such combinations should be experimentally verified as there can be unexpected negative effects when combining different NLS sequences^29^ (**Figure 2**). The improvement by the different NLSs over the SV40 NLS is probably the result of improved Cas9 nuclear targeting as the SV40 NLS results in GFP accumulation in both the nucleus and the cytoplasm (**Supplemental Figure 1b**). Unfortunately, we cannot directly test this as we have been unable to directly visualize Cas9 localization in plant cells as direct fluorescent protein fusions. Interestingly, the double-BP architecture was identified in optimization campaigns using Cas9 base editing in human cells and Cas12a-ABE in wheat^28, 33^, suggesting that this configuration may be generally applicable across a wide range of species and CRISPR systems. This would be particularly beneficial when using PAM-relaxed variants which have lower enzymatic activities than the wild-type versions^54^.

There is a large body of work testing promoters to express Cas9 in Arabidopsis (**Supplemental Table 1**). Numerous groups have identified ubiquitous promoters like RPS5A, UBQ1 and UBI10, and tissue-specific promoters such as YAO and EC1, however only two have compared these promoters side-by-side^2, 4^. These initial studies were limited because comparisons were made between plants with active Cas9 systems, and therefore somatic and germline editing cannot be distinguished. Our results show that this can be problematic when estimating the efficiency of a particular Cas9 promoter. For example, the RPS5A promoter only led to 64% of T1s with KO phenotypes (somatic and germline mutations) with GL1-2, as compared to PcUBI which gave 85% (**Figure 3b**). However, in T2 RPS5A had an average KO of 66% compared to PcUBI with an efficiency of 27% (**Figure 3c**). Genotyping at the Cas9-free T2 individuals is the best practice as one can be certain that the observed mutations are fixed in the genome, which is the overall goal of almost all mutagenesis projects.

Furthermore, only one or two target genes were used in these earlier reports, making it difficult to know if the results can be extrapolated to other targets or how they will perform with multiplexing. Here, we build on those reports by demonstrating that the RPS5A promoter is indeed the most efficient at producing germline mutations when targeting up to seven genes at a time. While the HMG promoter can be highly efficient, it performs inconsistently across different target sites. This is an interesting observation, but not useful for the production of mutants. As such, we do not recommend using this promoter to produce KO lines. We were particularly surprised to find that the EC1 promoter was inefficient in our hands, considering its widespread use in the field. Likewise, the YAO promoter was found to be an efficient promoter in multiple comparative studies, but failed to produce a high number of KO T2 seedlings in our system^2, 3, 12^. These discrepancies may be due to our use of Cas9-free T2s, our vector assembly (backbone, orientation of expression cassettes, etc.) or target-specific effects as we observed with the HMG promoter. Considering that we and others have repeatedly identified the RPS5A promoter^2, 4, 5^, we think that this is currently the most reliable promoter to use in Arabidopsis for the production of KO lines.

With these optimizations there is, of course, still room for improvement. While we have shifted the distribution to 4-6-plex mutants, even higher levels will be needed to rapidly mutate genomes and large gene families. It would be interesting to combine the RPS5A::BP-BP vector with different strategies to express gRNAs and/or by including introns in the Cas9 coding sequencing^32^. Experimenting with the different gRNA expression strategies would be especially interesting given that the gRNAs are the limiting factor in the RPS5A::BP-BP vector. This may be an inherent feature of certain gRNA sequences or it could be the result of the production of the RNA sequence *in vivo*. Extensive work is also needed to evaluate the off-targeting rate using such high-efficiency vectors. Here, we only had a single gRNA that was suitable to evaluate for off-targeting. More systematic studies are needed to test if there is indeed a trade-off between activity and specificity.

A common practical question when starting on a new CRISPR experiment in plants is the number of plant lines that need to be generated and propagated to find a certain number of unique, multiplex mutant lines. In our experiments, we selected only 8-10 T1 lines for evaluation and with the PcUBI::-SV40 architecture, typically only two or three lines contained KO mutations. Thus, when using this architecture, researchers would probably want to produce at least 30 independent lines to ensure they obtain a few useful lines. By switching to the RPS5A promoter or double-BP architecture, we find nearly every single T1 line produced T2 seedlings with KO mutations. Therefore, relatively small populations of T1s (10-20) should be sufficient for standard CRISPR/Cas9 experiments targeting seven or fewer genes. At these efficiencies, it is not necessary to pre-screen lines by genotyping at the T1 generation. Lines can be rapidly screened at T2 with just a handful of individuals (∼8) to identify those lines with the highest-order KOs. More individuals from the best lines could then be screened for higher-order mutants.

Overall, the combination of the double-BP architecture and RPS5A components led to an average of 78% of alleles edited across all targets, effectively all T2 plants contained at least one inheritable KO mutation and the majority of plants contained 4-7-plex mutations. We expect this configuration will simplify the generation of transgene-free Arabidopsis mutant lines and would be applicable for mutagenesis in other plants or organisms.

## MATERIALS AND METHODS

### Vector construction

See **Supplemental Table 4** for a detailed breakdown of the different plasmids and how they were assembled.

Golden Gate gRNA entry vectors were constructed as previously described^42^. Briefly, DNA oligos were annealed and inserted into the gRNA entry vectors (containing an Arabidopsis U6-26 pol III promoter, a ccdB-CmR selection cassette, and a Cas9 gRNA scaffold) using a Golden Gate reaction with BbsI-HF (New England Biolabs). Gibson assembly reactions were done using 2x NEBuilder Hifi DNA Assembly Mix (New England Biolabs) and incubated at 50 °C for 15 minutes. All gRNA sequences can be found in **Supplemental Table 5**.

Golden Gate destination and expression vectors were constructed as previously described^42^ or via an in-house “golden Gibson” protocol. Briefly, equimolar concentrations of the vectors were mixed with 1 µL I-SceI restriction enzyme (New England Biolabs) and 1 µL CutSmart buffer (10x) in a total volume of 10 µL. This reaction was incubated for two hours at 37°C and 25 minutes at 65°C. Five µL of the reaction was mixed with 5 µL 2x NEBuilder Hifi DNA Assembly Mix and incubated for one hour at 50°C.

Cloning reactions were transformed into One Shot™ ccdB Survival™ 2 T1R Competent Cells (Thermo Fisher Scientific) or ccdB-sensitive DH5α *Escherichia coli* via heat-shock. *E. coli* were cultivated on lysogeny broth medium containing 100 µg/mL spectinomycin or carbenicillin. Colonies were verified via colony-touch PCR, restriction digestion, Sanger sequencing (Eurofins Scientific), and/or whole plasmid sequencing (SNPsaurus or Eurofins Scientific).

We modelled the double-BP architecture on the BE4max and ABE8e configuration for base editing^28, 55^ where the C-terminal BP NLS lacks a serine as compared the N-terminal BP NLS. All NLS sequences can be found in **Supplemental Table 6**.

### Agrobacterium transformation and Nicotiana benthamiana agroinfiltration

Plant transformation vectors were transformed into *Agrobacterium tumefaciens* C58C1 by electroporation. Colonies were checked via colony PCR. *Agrobacterium* clones were grown overnight in YEB medium supplemented with 10 mM MES, 20 µM acetosyringone and appropriate antibiotics. Bacterial cells were pelleted, resuspended in infiltration medium (10 mM MgCl_2_, 10 mM MES, 100 µM acetosyringone) to an OD_600_ of 1 and incubated for 2-3 hours in a shaker at 28° and 150 rpm. The expression-vector cultures were mixed with a P19 culture in a 1:1 ratio. The abaxial side of the *N. benthamiana* leaf was punctured with a needle and the Agrobacterium mixture infiltrated using a 1 mL TERUMO®SYRINGE without a needle. Leaves were imaged three days after infiltration.

### PSB-D transformation and protoplasting

*Arabidopsis thaliana* cell suspension cultures (ecotype Landsberg *erecta*; PSB-D), derived from MM2d^56^, were maintained and transformed as previously described^57^. Transformation of the mutagenesis experiments targeting *GL1* was performed with a single *Agrobacterium* clone. Transformation of the multiplex mutagenesis vectors was performed with an *Agrobacterium* solution containing three clones that were mixed at the *Agrobacterium* inoculation step. The cell suspension culture was protoplasted 17 days post-cocultivation as previously described^45^. Protoplasts were sorted on the BD FACSMelody (BD Biosciences) equipped with three lasers (405 nm, 488 nm, and 561 nm). The gating strategy for each experiment is shown in **Supplemental Figure 14**. In total, 75,000 mCherry-positive protoplasts were collected in Mannitol magnesium solution and maintained on ice. The samples were centrifuged for 2 minutes at 4000 rpm and the supernatant was removed before storage at −80°C.

### Plant transformation and DNA extraction

Transformation of *Arabidopsis thaliana* Col-0 plants was performed via the floral-dip method^41^. FASTR-positive or -negative seeds were selected and grown as previously described^42^. Seeds were sown on half strength MS medium containing 200 mg/L timentin and/or 50 mg/L kanamycin. After stratification of two days, the seeds were transferred to a growth chamber (21 °C, 16h light and 8h dark regime). After two weeks, the first true leaves were harvested, the seedlings were transferred to Jiffy-7 pellets, and grown in a greenhouse under a 16h light and 8h dark regime. DNA was extracted according to either Berendzen et al. (2005) or Edwards et al. (1991) with modifications as previously described (Develtere et al., 2023).

### DNA sequencing

For Sanger sequencing of the PSB-D protoplasts, target regions were amplified using the ALLin^TM^ Red Taq Master Mix (highQu) according to the manufacturer’s instructions. Bead-purified PCR amplicons (CleanNGS) were sequenced via the Mix2Seq service (Eurofins Genomics).

For Illumina amplicon sequencing, PCRs were performed with the iProof High-fidelity DNA polymerase according to the manufacturer’s instructions. Primers to amplify the target regions were barcoded with a 6-nt tag^58^ to allow pooling. All target regions were amplified in one reaction with a final primer concentration of 0.5 µM each. A maximum of 96 PCR reactions were pooled at a time and cleaned with Zymo-Spin II columns (Zymo Research). Pooled samples were sequenced with the NGSelect Amplicon sequencing service of Eurofins Genomics.

### DNA sequence analysis

Sanger sequencing results were analysed with the ICE software^46^. The Illumina reads were demultiplexed^59^, and the read pairs overlapped, merged^60^, and trimmed^61^. The reads were mapped to the reference using BWA-MEM^62^ (See **Supplemental File 1** for reference sequences). The mapped reads were analysed using SMAP haplotype-window^49^ with the default parameters except for *min. read count,* 30; *min. haplotype frequency*, 5. For quality control we plotted the frequency of each genotype score for each experiment (**Supplemental Figure 3a**). For the transformed lines we observed a trimodal pattern, with peaks at 0, 50, and 100%, as expected for a diploid. For the wild-type samples, we observe a peak between 0 and 10%. This allows us to make discrete genotype calls, with 20% being the threshold for calling an allele mutated. Per plant, a target site was called WT if the haplotype frequency of the reference sequence was > 80%. A target was called heterozygous if the reference haplotype and exactly one non-reference haplotype each had a frequency between 20% and 80%. A target was called homozygous if exactly one non-reference haplotype had a frequency >80%, and bi-allelic if exactly two distinct non-reference haplotypes each had a frequency between 20% and 80%. If none of these criteria were met, the genotype call for that target locus was called “false” and excluded from the analyses. For quality control, we plotted all calls per plant (**Supplemental Figure 4**). All genotype calls can be found in **Supplemental File 3**.

### Confocal microscopy

Infiltrated *N. benthamiana* leaves were imaged on an Olympus FluoView™ FV1000 confocal microscope. GFP was excited at 488 nm and acquired at 500 nm to 545 nm.

### Statistical Analysis

The “editing” data were treated as binary, taking no edit = 0, and edited = 1. Having several T2 plants derived from a single T1 plant and/or multiple targets within T2 plants subjected to the editing process, the data were collapsed into binomial data, with a total of n targets (either genes or alleles) and r being edited. For example, having 11 T2 plants/T1 plant for which a binary response has been recorded in 7 genes, the data were collapsed into a total of 11×2×7 = 154 target alleles of which r being edited. Because of the binomial nature of the data, a logistic regression model, with a logit link function, as implemented in Genstat (version 22, VSN International) was fitted to the response data. The dispersion parameter for the variance of the response was estimated from the residual mean square of the fitted model. Wald statistics were used to assess the significance of the main treatments and their interaction terms, by dropping these fixed terms from the full model. T-statistics were used to assess the significance of treatment (e.g. promotor, promotor-NLS, etc…) effects (on the logit transformed scale) by pairwise comparisons to reference treatment level. All statistical test results and input data can be found in Supplemental File 3.

The ICE scores were treated as continuous data and analyzed by a two-way ANOVA. The significance of the main terms M and NLS and their interaction was assessed by an F-test as implemented in Genstat (version 22, VSN International).

## DATA AVAILABILITY

MAS files can be found in the SRA database (https://www.ncbi.nlm.nih.gov/sra) under the accession number PRJNA1036854. All source data to generate graphs are available in Supplemental File 2 and Supplemental File 3. Plasmids and vector maps (.gb files) are available at https://gatewayvectors.vib.be/.

## ACKNOWLEDGMENTS

We thank Mansour Karimi for general cloning support and providing the Golden Gibson cloning vectors. We thank Carina Braeckman for performing the Arabidopsis floral-dip transformation. We thank Nico Smet, Miguel Riobello y Barea, Thomas Farla, and Sandra Lefftz for greenhouse support. We thank Carina Braeckman, Nancy Helderwert and Alexandra Peña Fernandez for help with harvesting. We thank Rafael Andrade Buono and Jonas Blomme for their help with confocal imaging. We thank Julie Van Duyse and Gert Van Isterdael from the VIB Flow core for assistance with FACS.

