## Supplemental Tables and Figures for "Continual improvement of multiplex mutagenesis in Arabidopsis"

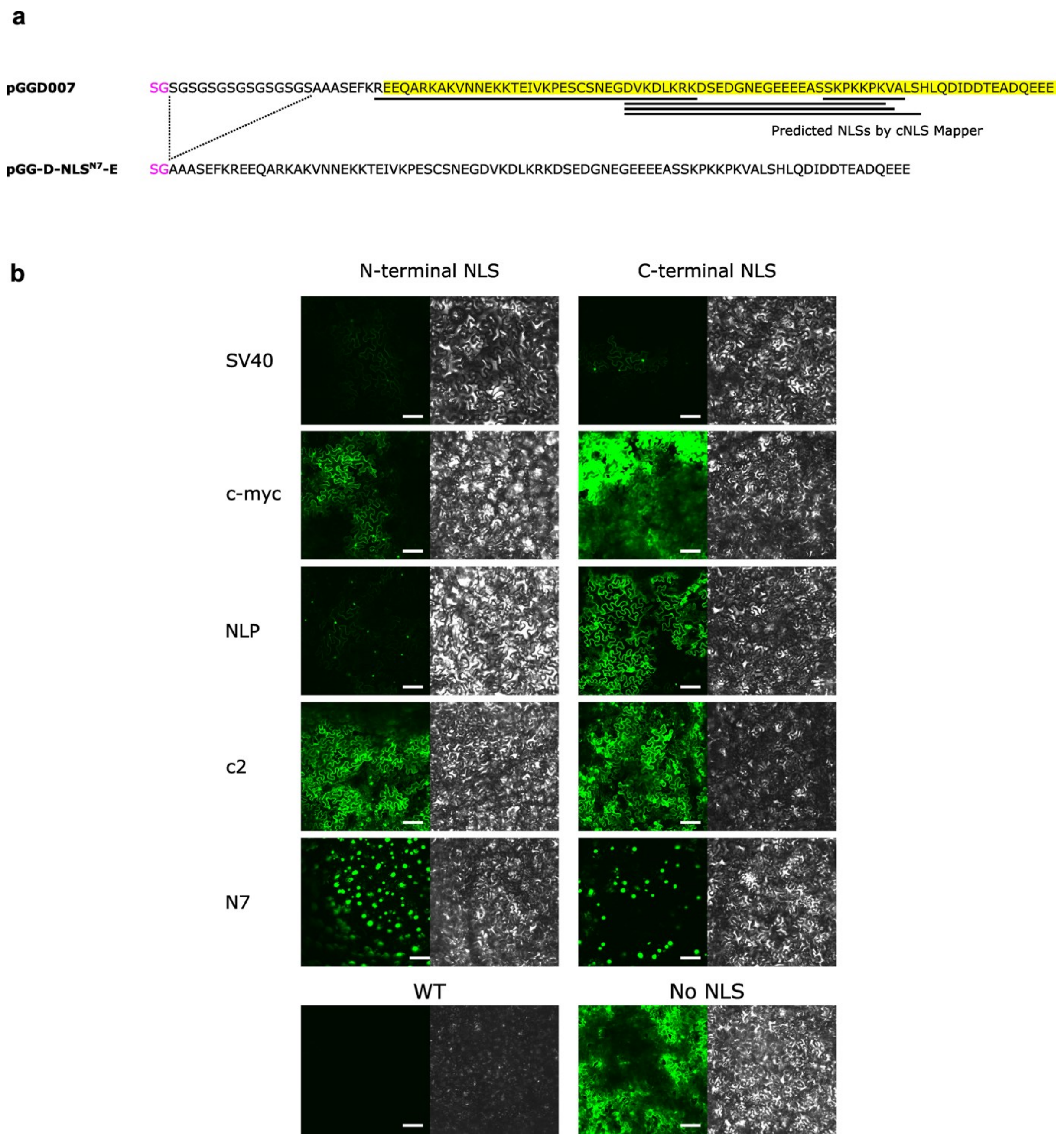


**Supplemental Figure 1: The N7 NLS and testing different NLS-GFP fusions *in planta*** (**a**) Amino acid sequence of the N7 cloning module and predicted NLSs. (top) Sequence of pGGD007 and (bottom) pGG-D-NLSN7-E. The sequence highlighted in yellow represents the C-terminal region of the Ankyrin repeat family protein (AT4G19150). The sequence in purple represents the cloning scar. Predicted NLSs by cNLS Mapper are underlined. (**b**) Agroinfiltration of *Nicotiana benthamiana* with different vectors expressing GFP-NLS fusion proteins. WT: Picture of a wild-type plant, No NLS: infiltration of control vector lacking an NLS sequence. One representative image was chosen per condition. The scale bar is 100 µm.

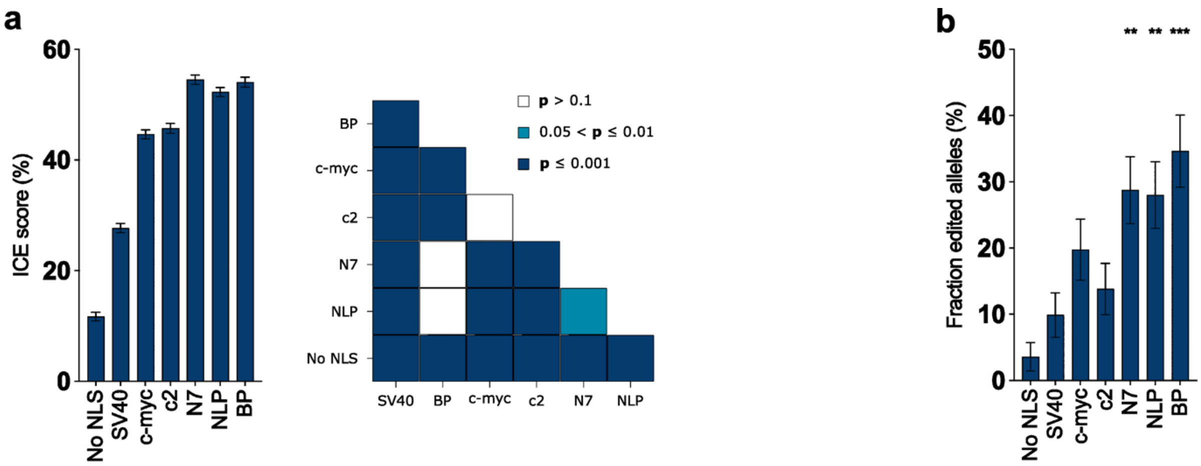


**Supplemental Figure 2: Efficiency comparison of different C-terminal NLS fusions.** (**a**) Vectors containing a single C-terminal NLS fusion were transfected into PSB-D cell cultures. The mean ICE score (from three or four replicates) across the 14 targets per vector (left) and the p-values between vectors (right) are shown. The error bars represent the standard error of the mean (two-way ANOVA followed by a Fisher's protected Least Significant Difference Test). (**b**) The same vectors as (a) were transformed into Arabidopsis plants. The mean fraction of edited alleles across the 14 targets per vector in the T2 generation is shown. The error bars represent the standard error of the mean. Comparisons of each architecture with the SV40 architecture were tested for significance (logistic regression followed by Student's two-sided t-testing of the pairwise differences of the estimates to the reference (on the logit scale)). **, $p < 0.01$; ***, $p < 0.001$. Exact p-values are shown in Supplementary File 2.

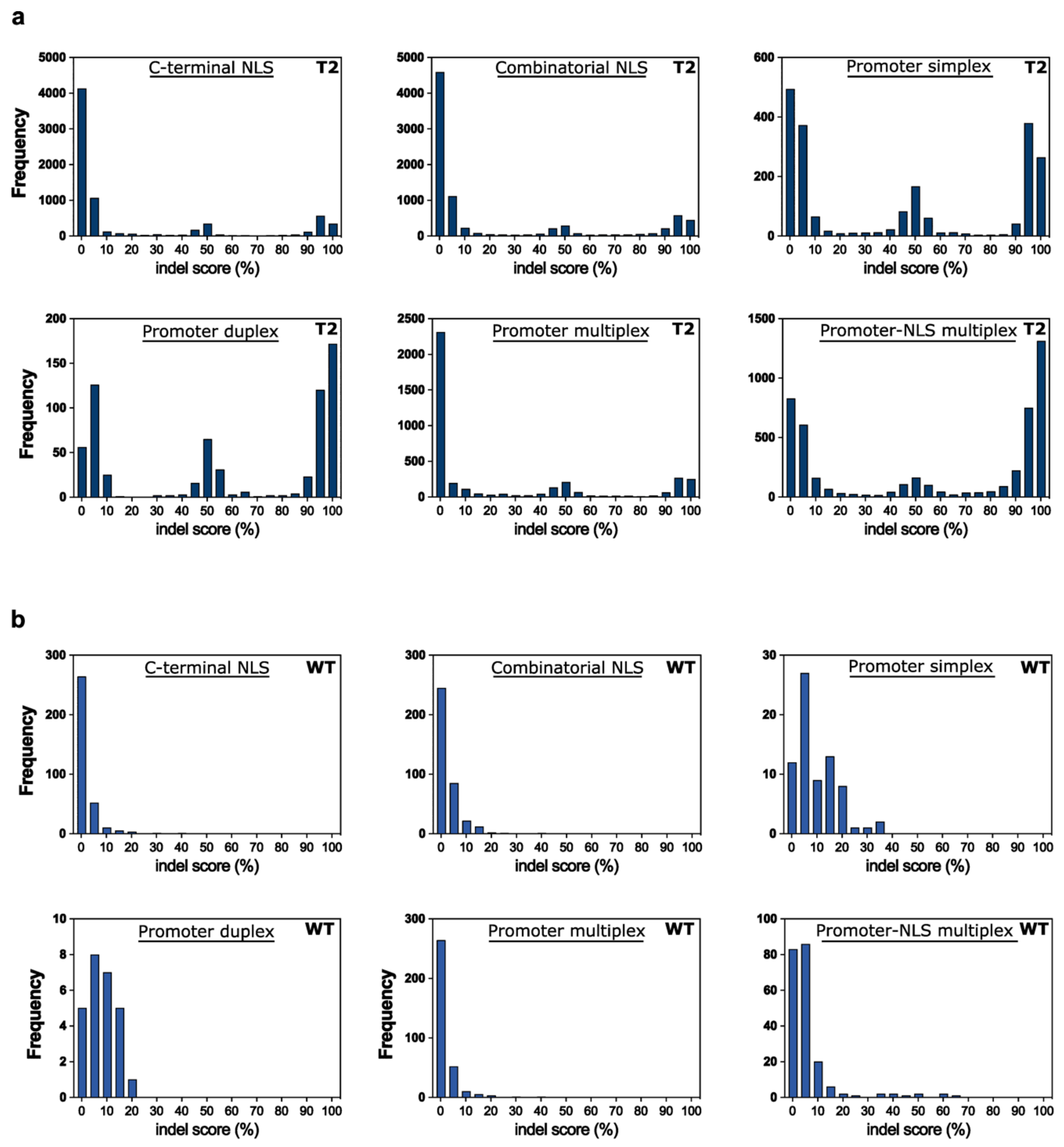


**Supplemental Figure 3: Indel scores of the different datasets.** For each multiplex amplicon sequencing dataset, the indel score (=100 – WT haplotype frequency) of each target of each T2 sample (**a**) and WT sample (**b**) was calculated and binned. The frequency of each bin per dataset is shown.

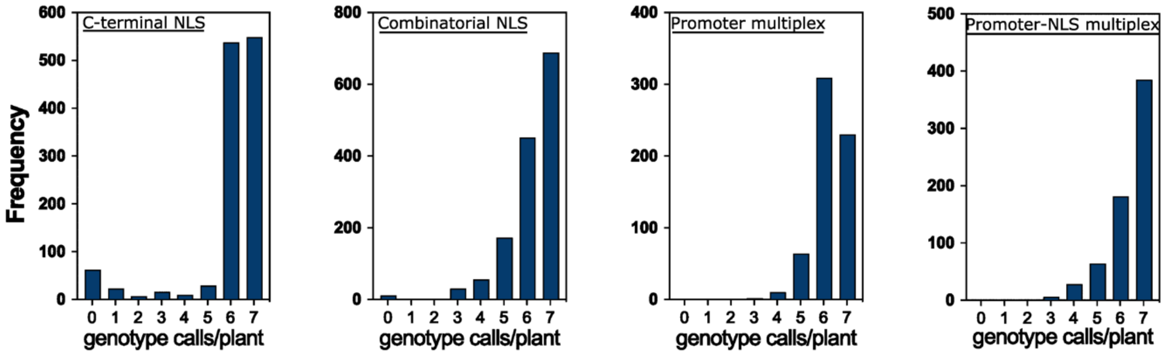


**Supplemental Figure 4: Genotyping coverage of the different datasets.** Seven target sites per plant were genotyped (M1 or M2) for the different multiplex comparison experiments using multiplex amplicon sequencing. Genotype calls were given for each target site for each plant based on the read frequency. The frequency of the number of genotype calls per plant is given for each experiment.

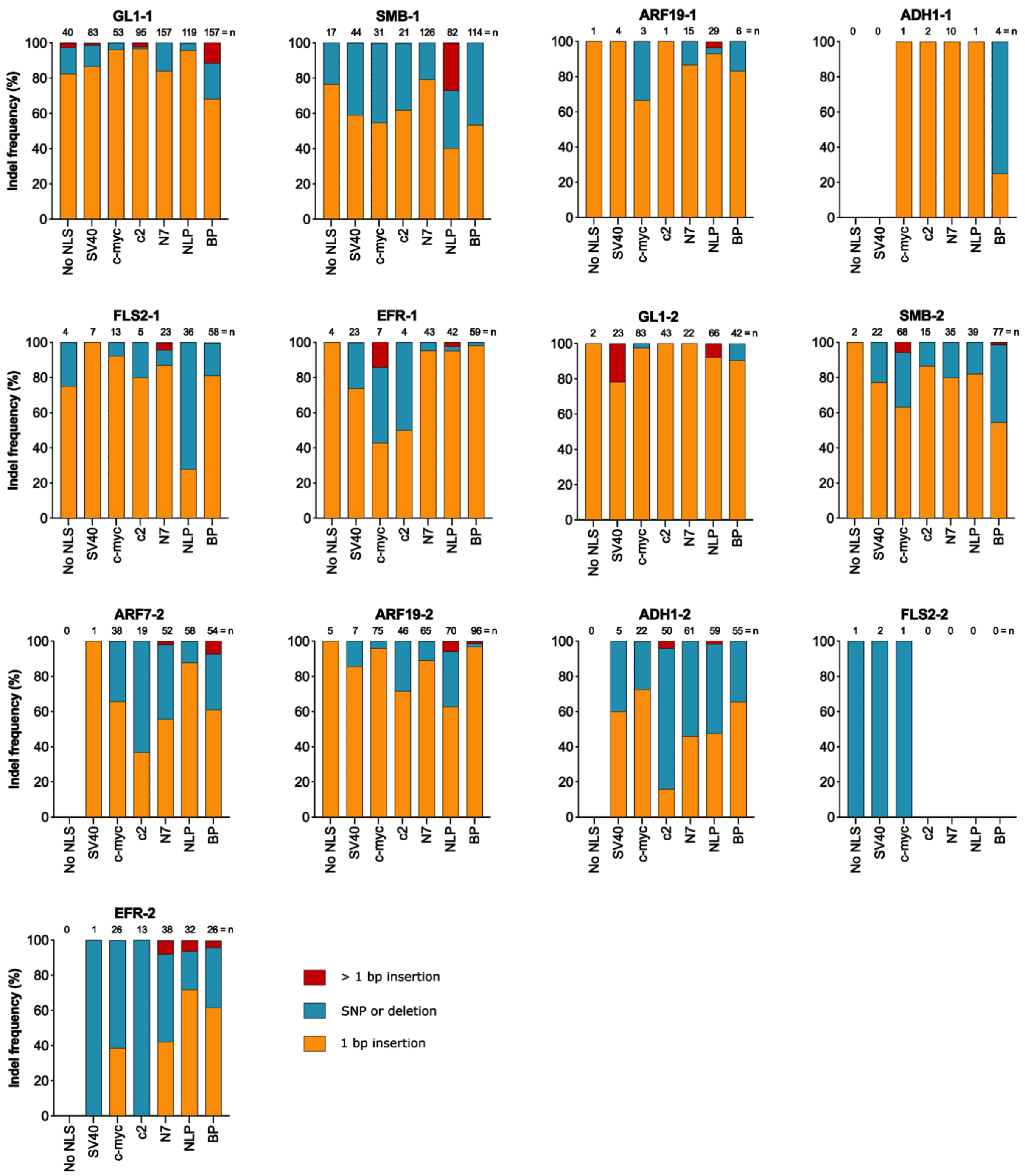


**Supplemental Figure 5: Indel size at different target sites of the C-terminal NLS dataset.** The indel size at different edited target sites was grouped in either a 1 bp insertion, a SNP or deletion, or an insertion greater than 1 bp category. The counts were binned for the different architectures. The frequency of each group is given. The MAS for target ARF7-1 failed in this experiment. The sample size is given above the bars.

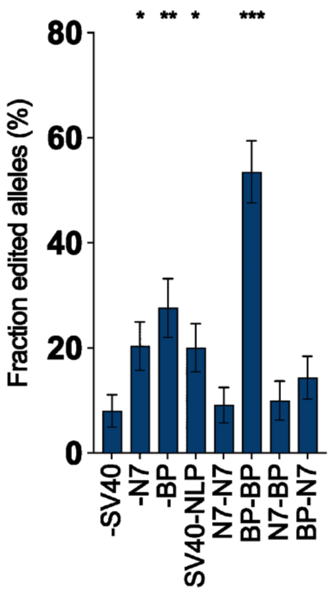


**Supplemental Figure 6: Efficiency comparison of different combinatorial NLS fusions**. Vectors containing a combinatorial NLS architecture and C-terminal-only NLS architectures were transformed into Arabidopsis plants. The mean fraction of edited alleles across the 14 targets per vector is given. The error bars represent the standard error of the mean. Comparisons of each architecture with the -SV40 architecture were tested for significance (logistic regression followed by Student's two-sided t-testing of the pairwise differences of the estimates to the reference (on the logit scale)). *, $p < 0.05$; **, $p < 0.01$; ***, $p < 0.001$. Exact p-values are shown in Supplementary File 2.

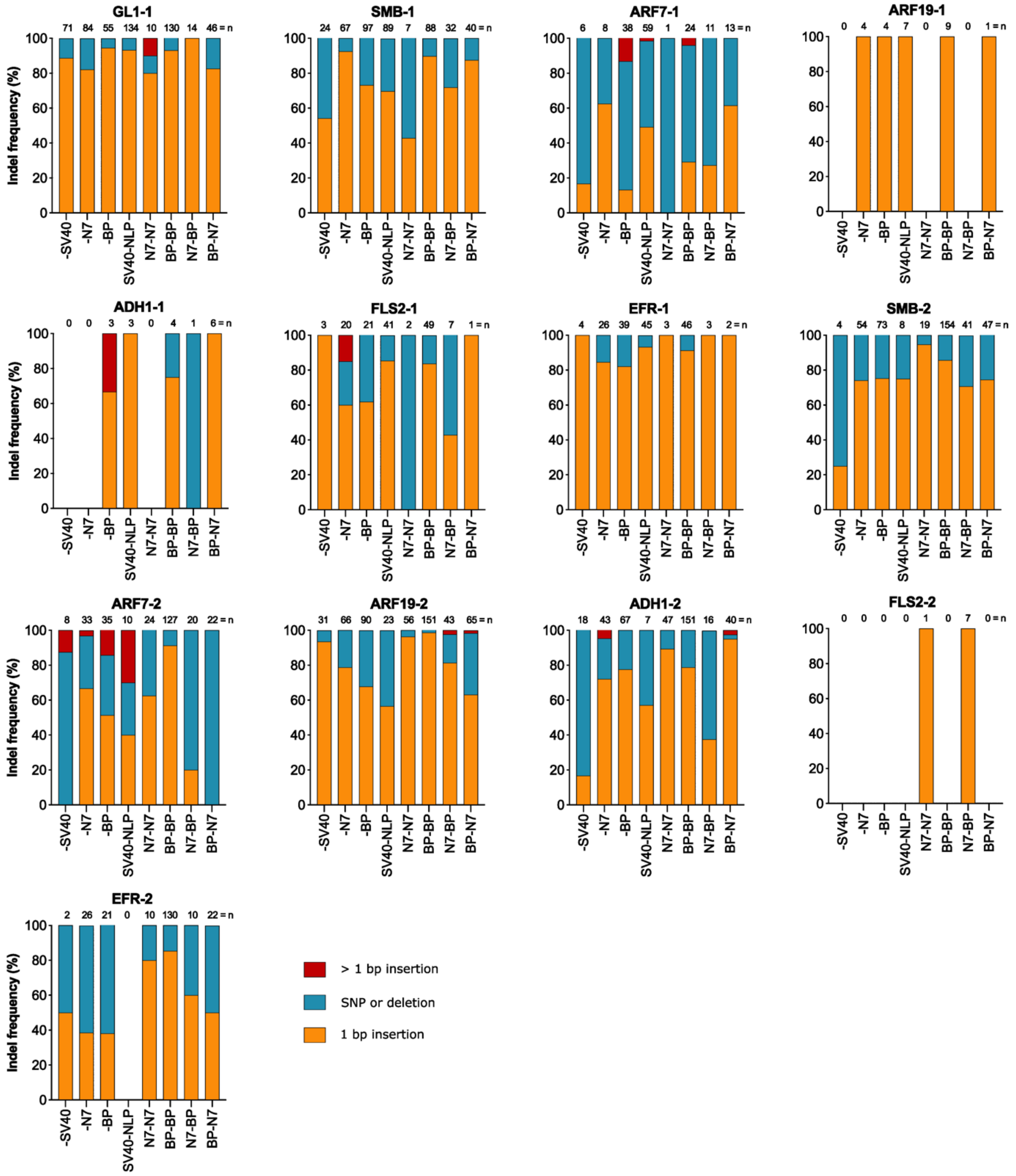


**Supplemental Figure 7: Indel size at different target sites of the combinatorial NLS dataset.** The indel size at different edited target sites was grouped in either a 1 bp insertion, a SNP or deletion, or an insertion greater than 1 bp category. The counts were binned for the different architectures. The frequency of each group is given. The multiplex amplicon sequencing for target GL1-2 failed in this experiment. The sample size is given above the bars.

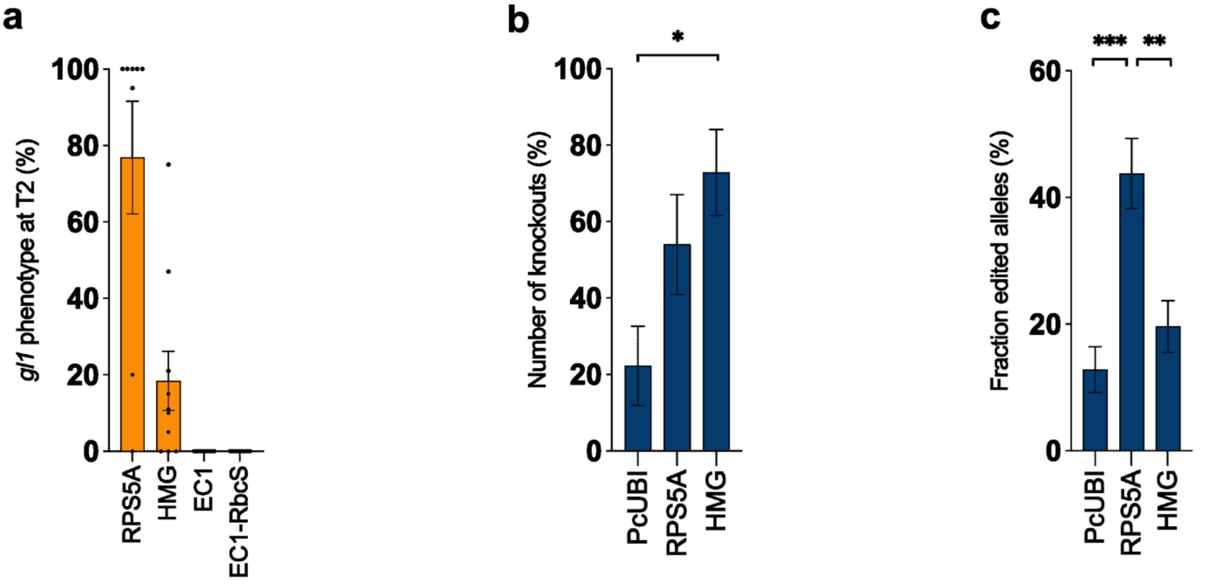


**Supplemental Figure 8: Efficiency comparison of different promoters expressing Cas9**. (**a**) The efficiency of the RPS5A, HMG and EC1 vectors with G7 terminator were compared to the EC1 vector with RbcS terminator with the GL1-2 gRNA. 5 – 10 T1 lines were propagated to T2, and for each T1 line 16 – 21 T2 seedlings were screened for the KO phenotype. (**b**) The percentage of all possible knockouts across all lines per promoter with the ARF7-2 and ARF19-2 gRNAs in duplex is given. The error bars represent the standard error of the mean (logistic regression followed by Student's two-sided t-testing of the pairwise differences of the estimates to the reference (on the logit scale)). (**c**) The mean fraction of edited alleles across the 14 targets per vector is given. The error bars represent the standard error of the mean (logistic regression followed by Student's two-sided t-testing of the pairwise differences of the estimates to the reference (on the logit scale)). *, $p < 0.05$; **, $p < 0.01$; ***, $p < 0.001$. Exact p-values are shown in Supplementary File 2.

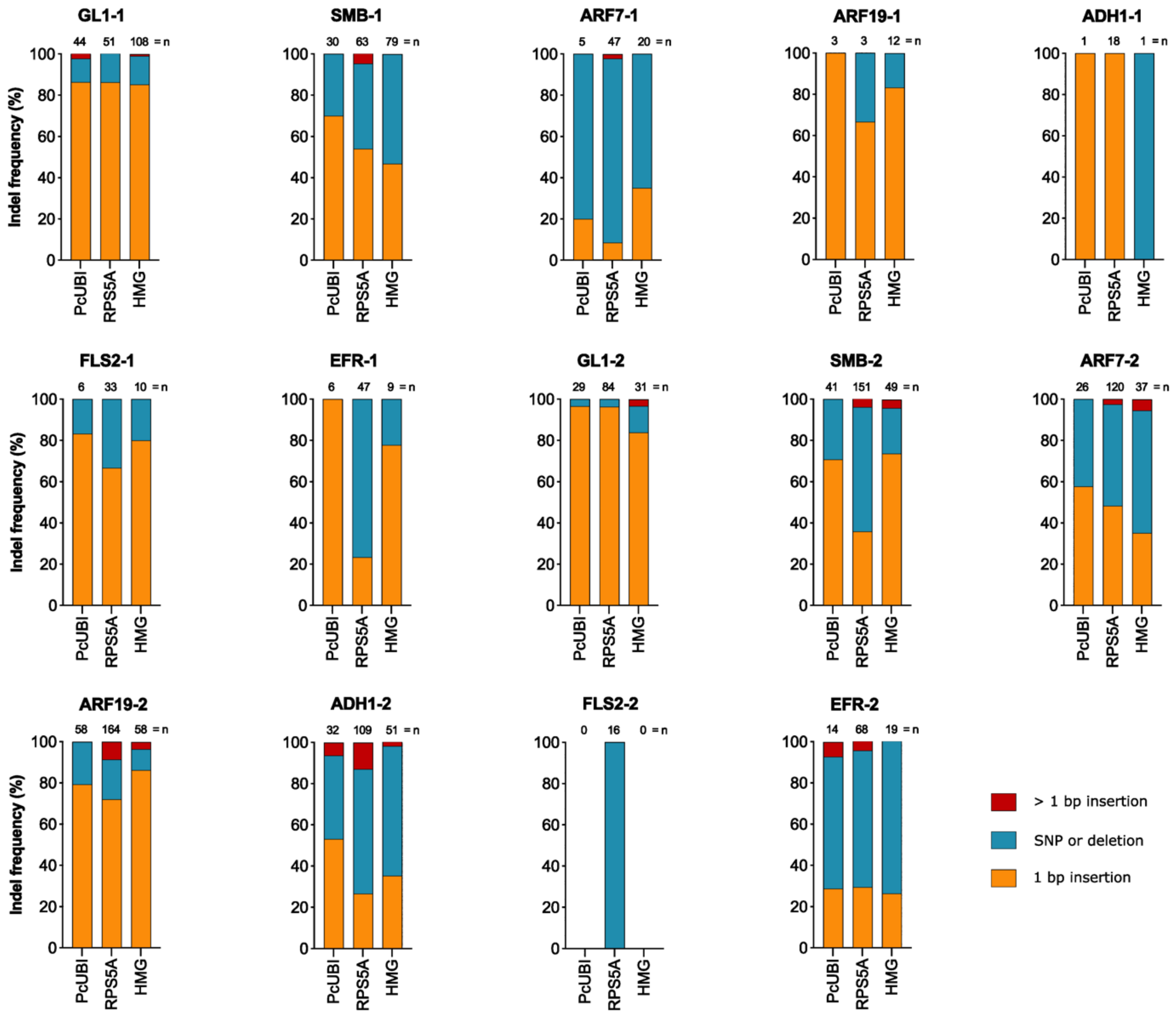


**Supplemental Figure 9: Indel size at different target sites of the promoter multiplex dataset.** The indel size at different edited target sites was grouped in either a 1 bp insertion, a SNP or deletion, or an insertion greater than 1 bp category. The counts were binned for the different promoters. The frequency of each group is given. The sample size is given above the bars.

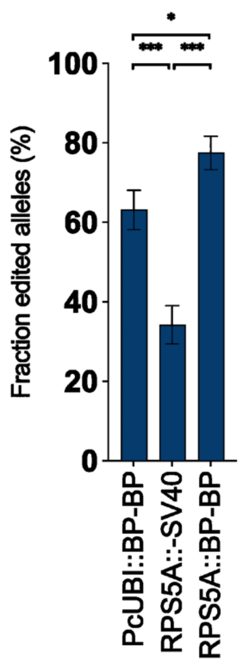


**Supplemental Figure 10: Efficiency comparison of different promoters and NLS combinations expressing Cas9**. The best performing promoter and NLS were combined and transformed into Arabidopsis alongside the best performing architectures found in previous experiments. The mean fraction of edited alleles across the 14 targets per vector is given. The error bars represent the standard error of the mean (logistic regression followed by Student's two-sided t-testing of the pairwise differences of the estimates to the reference (on the logit scale)). *, $p < 0.05$; ***, $p < 0.001$. Exact p-values are shown in Supplementary File 2.

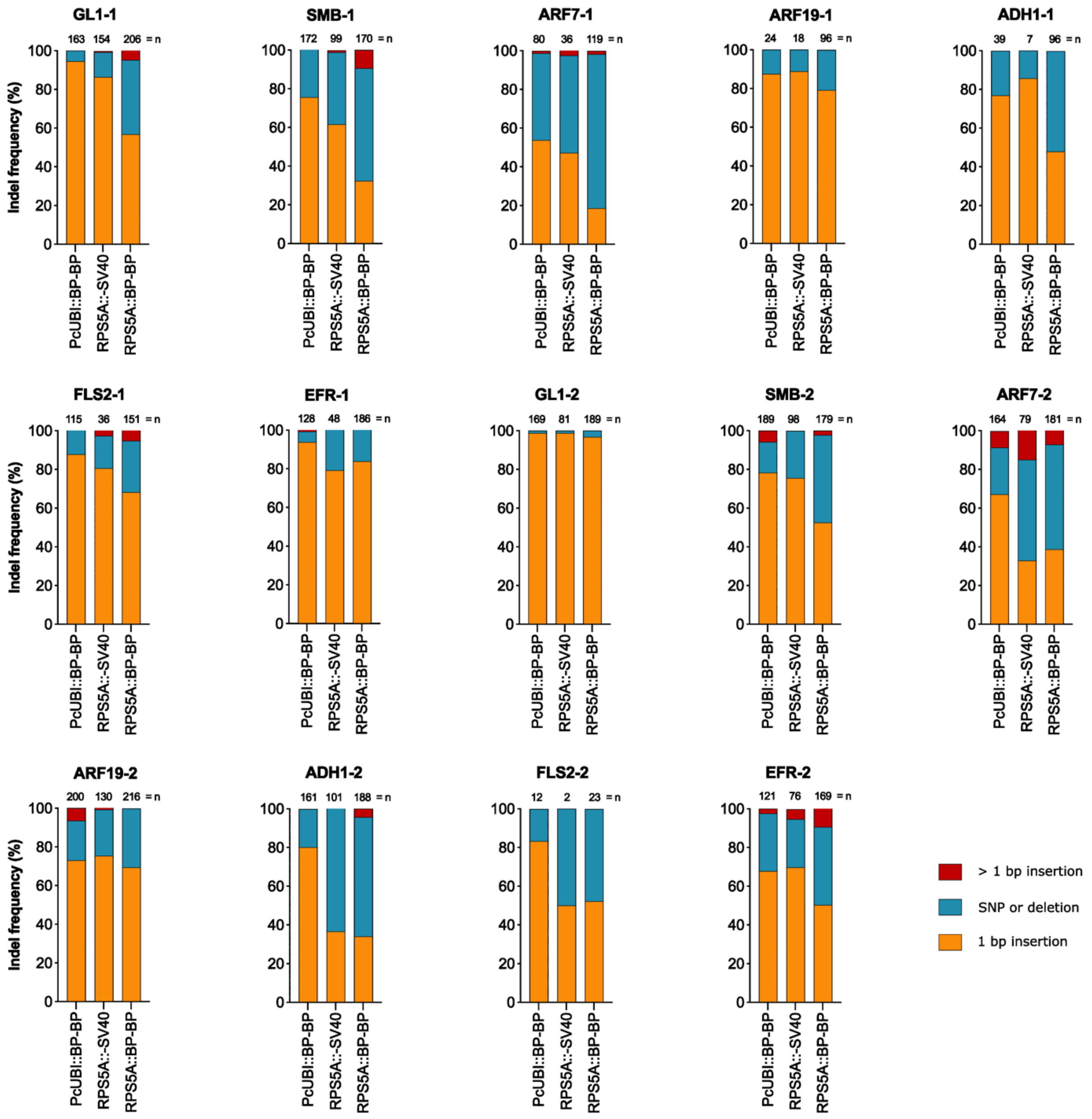


**Supplemental Figure 11: Indel size at different target sites of the promoter-NLS multiplex dataset.** The indel size at different edited target sites was grouped in either a 1 bp insertion, a SNP or deletion, or an insertion greater than 1 bp category. The counts were binned for the different promoters. The frequency of each group is given. The sample size is given above the bars.

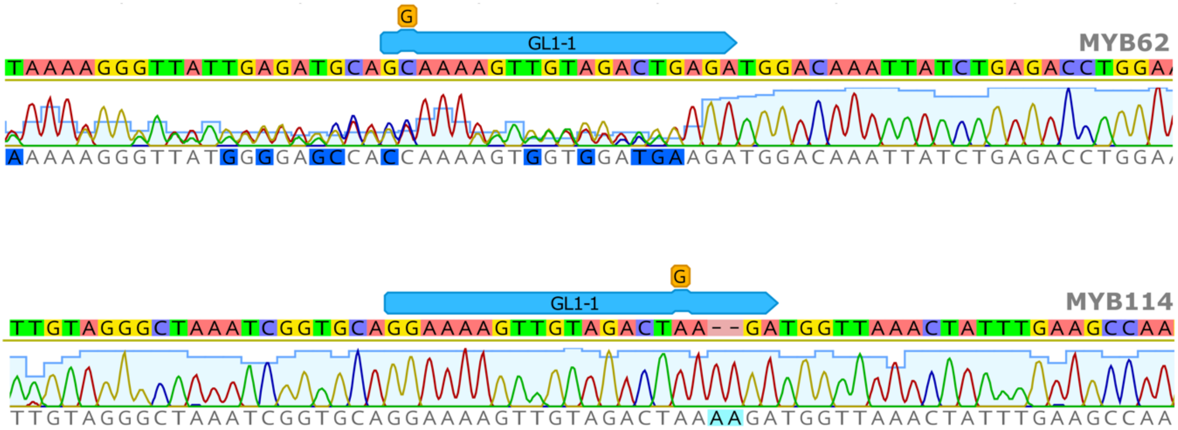


**Supplemental Figure 12: Sanger sequencing of GL1-1 off-target loci**. Four potential off-target sites for GL1-1 were sequenced to assess the off-target activity. One off-target event was identified in gene *MYB62* (top panel) and one in gene *MYB114* (bottom panel). The chromatogram and base-calls for both events are shown. The gRNA is shown in blue. The mismatch of the gRNA with the reference is shown as an orange rectangle.

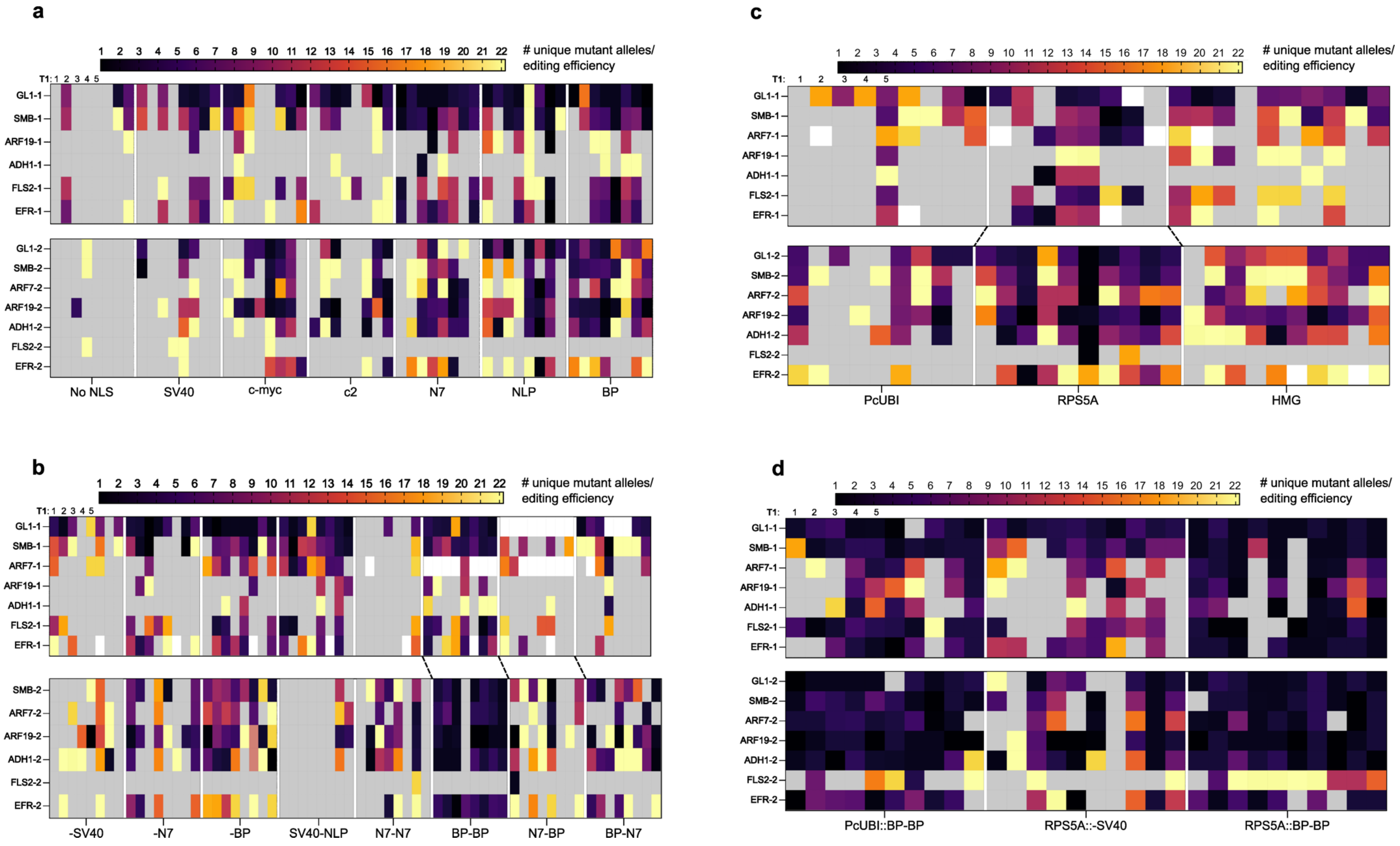
a
b
c
d
### unique mutant alleles/ editing efficiency
T1: 1 2 3 4 5
GL1-1
SMB-1
ARF7-1
ARF19-1
ADH1-1
FLS2-1
EFR-1
GL1-2
SMB-2
ARF7-2
ARF19-2
ADH1-2
FLS2-2
EFR-2
No NLS
SV40
c-myc
c2
N7
NLP
BP
-SV40
-N7
-BP
SV40-NLP
N7-N7
BP-BP
N7-BP
BP-N7
PcUBI
RPS5A
HMG
PcUBI::BP-BP
RPS5A::-SV40
RPS5A::BP-BP

**Supplemental Figure 13: Diversity of alleles of the different datasets.** The allele diversity was calculated for the C-terminal NLS (**a**), Combinatorial NLS (**b**), Promoter multiplex (**c**), and Promoter-NLS multiplex (**d)** datasets. Per T1 line, the number of unique mutant alleles was divided by the editing efficiency of that line and target. White cells indicate missing data.

a

b

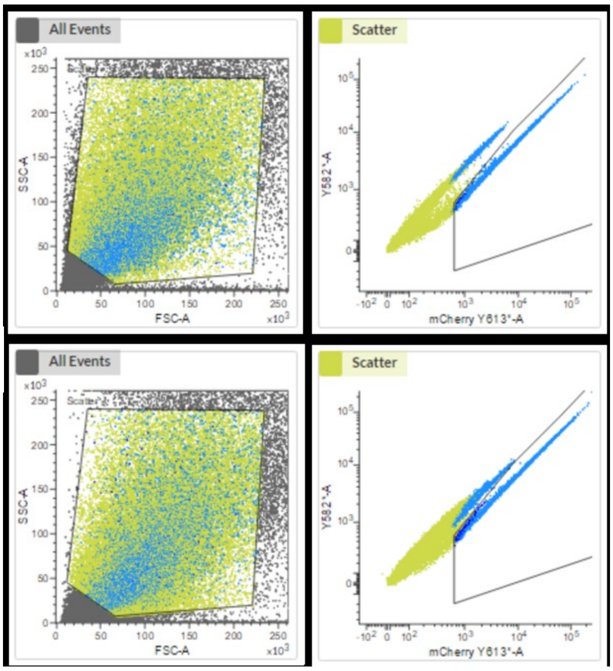


**Supplemental Figure 14: FACS gating settings for sorting mCherry-positive protoplasts.** Vectors containing a single C-terminal NLS fusion were transfected into PSB-D cell cultures. The cultures were protoplasted and sorted with FACS. The gating settings for (**a**) M1 and (**b**) M2 are shown.

**Supplemental Table 2. Most popular CRISPR plasmids on Addgene in plants (as of November 22, 2023)**. Linker sequences are underlined.

| Rank | Plasmid # | Name | Architecture | Depositing Lab | Reference |
|---|---|---|---|---|---|
| 1 | 160393 | JD633 | M-NLS_**SV40**-GIHGVPAA-Cas9-NLS_**NLP** | Jorge Dubcovsky | Debernardi et al 2020 |
| 2 | 63142 | pRGEB32 | 3xFLAG-MA-NLS_**SV40**-GIHGVPAA-Cas9-NLS_**NLP** | Yinong Yang | Xie et al., 2015 |
| 3 | 62201 | pHSE401 | 3xFLAG-MA-NLS_**SV40**-GIHGVPAA-Cas9-NLS_**NLP** | Qi-Jun Chen | Xing et al., 2014 |
| 4 | 71287 | pHEE401E | 3xFLAG-MA-NLS_**SV40**-GIHGVPAA-Cas9-NLS_**NLP** | Qi-Jun Chen | Wang et al., 2015 |
| 5 | 91135 | pDIRECT_22C | Csy4-P2A-Cas9-SRAD-NLS_**SV40** | Daniel Voytas | Cermak et al., 2017 |
| 6 | 51295 | pRGEB31 | 3xFLAG-MA-NLS_**SV40**-GIHGVPAA-Cas9-NLS_**NLP** | Yinong Yang | Xie and Yang, 2013 |
| 7 | 62202 | pKSE401 | 3xFLAG-MA-NLS_**SV40**-GIHGVPAA-Cas9-NLS_**NLP** | Qi-Jun Chen | Xing et al., 2014 |
| 8 | 85758 | pKIR1.1 | 3xFLAG-MA-NLS_**SV4**0-GIHGVPAA-Cas9-NLS_**NLP** | Tetsuya Higashiyama | Tsutsui and Higashiyama, 2017 |
| 9 | 86210 | pYPQ230 (LbCpf1) | MA-NLS_**SV40**-GIHGVPAA-LbCas12a-NLS_**NLP** | Yiping Qi | Tang et al., 2017 |
| 10 | 85808 | pKI1.1R | 3xFLAG-MA-NLS_**SV40**-GIHGVPAA-Cas9-NLS_**NLP** | Tetsuya Higashiyama | Tsutsui and Higashiyama, 2017 |
| 11 | 52256 | pFGC-pcoCas9 | 2xFLAG-MA-NLS_**SV40**-GIHGVPAA-Cas9 (intron)-NLS_**NLP** | Jen Sheen | unpublished |
| 12 | 46965 | pK7WGF2::hCas9 | EGFP-DITSLYKKAGSAAAPFT-Cas9-SRAD-NLS_**SV40** | Sophien Kamoun | Nekrasov et al., 2013 |
| 13 | 52254 | HBT-pcoCas9 | 2xFLAG-MA-NLS_**SV40**-GIHGVPAA-Cas9 (intron)-NLS_**NLP** | Jen Sheen | Li et al., 2013 |
| 14 | 62203 | pHUE411 | 3xFLAG-MA-NLS_**SV40**-GIHGVPAA-Cas9-NLS_**NLP** | Qi-Jun Chen | Xing et al., 2014 |
| | 66187 | pYLCRISPR/Cas9Pubi-H | MA-NLS_**SV40**-GIHGVPAA-Cas9-NLS_**NLP** | Yao-Guang Liu | Ma et al., 2015 |
| | 49771 | pICH47742:2x35S-5'UTR-hCas9(STOP)-NOST | Cas9-SRAD-NLS_**SV40** | Sophien Kamoun | Belhaj et al., 2013 |
| | 153210 | pAGM55261 | MASSP-NLS_**SV40**-SWKM-Cas9-SRAD-NLS_**SV40** | Sylvestre Marillonnet | Grutzner et al., 2020 |

Accessed November 22, 2023

**Supplemental Table 3: Genotyping results confirm the mutation in knockout plants.** Up to four transgene-free T2 plants without trichomes from the transformation with GFP-Cas9-SV40 and GFP-Cas9-N7 were genotyped at the GL1-1 and GL1-2 loci with sanger sequencing. '+', insertion; '-', deletion.

| | **NLS** | **T1 Line** | **T1 Trichomes** | **T2 without Trichomes** | **Plant #1** | **Plant #2** | **Plant #3** | **Plant #4** |
|---|---|---|---|---|---|---|---|---|
| GL1-1 | SV40 | # 2 | No | 13/60 | +C | +C | +C | +C |
| | | # 9 | Yes | 23/38 | +T | +A/+T | | |
| | | # 13 | No | 44/63 | +A/+T | +A/-1 | +A | |
| | N7 | # 6 | No | 58/58 | +A/+T | +A/-9 | +A/-9 | +A/-9 |
| | | # 13 | No | 51/51 | -1/-6 | -6 | -6 | +T |
| | | # 18 | No | 42/52 | +A/+T | +A | +A | +A |
| GL1-2 | SV40 | # 6 | Yes | 3/60 | +A/-19 | +A/-14 | +T | |
| | | # 12 | Yes | 6/50 | +A/+T | +A/+T | +A/+G | -1 |
| | N7 | # 5 | No | 31/49 | +A | +T | +A/+G | +A/+G |
| | | # 9 | No | 44/45 | +A/+T | +A | +A/+T | |
| | | # 17 | No | 22/57 | +T | +T | -1 | |

**Supplemental Table 5.** gRNA target sequences. PAM sequences are indicated in bold.

| gRNA | Target Sequence | Reference |
|---|---|---|
| GL1-1 | GGAAAAGTTGTAGACTGAGA**TGG** | Hahn et al., 2017 |
| GL1-2 | TTGAGCCCTAATGTGAACAA**AGG** | |
| SMB-1 | GACTGGGACCCGAACGAACC**GGG** | Decaestecker et al., 2019 |
| SMB-2 | TACACTGGTCGTGCTCCTCA**TGG** | Decaestecker et al., 2019 |
| ARF7-1 | GCGCTGCACAACAATCTTGG**CGG** | Decaestecker et al., 2019 |
| ARF7-2 | GCGAGTGACACAAGTACTCA**CGG** | Decaestecker et al., 2019 |
| ARF19-1 | GGCCAAGCCGAGTATCCATA**TGG** | Decaestecker et al., 2019 |
| ARF19-2 | GAGAAATCACAGCTGATGTT**GGG** | Decaestecker et al., 2019 |
| ADH1-1 | CTCAGGATCAACACCGAGCG**AGG** | |
| ADH1-2 | TCAGGTTGCTAAGATCAATC**CGG** | |
| FLS2-1 | GTGAGCTCCCGGCGGATCTA**GGG** | |
| FLS2-2 | CTGCACCGGTCTTAAACTCC**TGG** | |
| EFR-1 | CCCTGAGAATAAACTCGTCG**CGG** | |
| EFR-2 | GTGGCTGCAGCTAGAAGATC**TGG** | |

**Supplemental Table 6.** NLS sequences used in this report. Additional amino acids, which are not part of the originally reported sequence, are underlined.

| NLS | Sequence |
|---|---|
| SV40 | SRADPKKKRKV |
| c-myc | PAAKRVKLD |
| NLP | KRPAATKKAGQAKKKKLD |
| c2 | QPSLKRMKIEPSSQP |
| N7 | AAASEFKREEQARKAKVNNEKKTEIVKPESCSNEGDVKDLKRKDSEDGNEGEEEEASSKPKKPKVALSHLQDIDDTEADQEEE |
| BP | KRTADGSEFEPKKKRKV |
